# Permute-match tests detect significant correlations between time series despite nonstationarity and limited replicates

**DOI:** 10.1101/2023.03.13.531689

**Authors:** Alex E. Yuan, Wenying Shou

**Affiliations:** Molecular and Cellular Biology PhD program, University of Washington and Fred Hutchinson Cancer Center, Seattle, United States; Centre for Life’s Origins and Evolution, Department of Genetics, Evolution and Environment, University College London, London, United Kingdom

## Abstract

Researchers frequently analyze correlations between pairs of time series by determining whether an observed correlation is stronger than expected under the null hypothesis of independence. However, the time series are often nonstationary, with statistical properties that change over time, thereby making standard tests invalid. If sufficient replicates exist, a trial-swapping permutation test can be performed that handles nonstationarity by comparing within-replicate correlations to between-replicate correlations. Although largely assumption-free, this test is fundamentally limited by the number of replicates (*n*) because its minimum p-value is 1*/n*!. With *n* = 3, this minimum is 1*/*6, rendering thresholds like 0.05 unattainable. This limits its use considerably in animal experiments, where *n* may be as low as 3. We propose permute-match tests — modified permutation tests that can report lower p-values of 2*/n*^*n*^ or 1*/n*^*n*^ under strong evidence of dependence. Permute-match tests guarantee a false positive rate at or below the significance level when replicates are independent and identically distributed. The bound of 1*/n*^*n*^ is not gratuitously conservative, since it cannot be further lowered without additional assumptions. We demonstrate our approach using synthetic data and apply it to an existing dataset with 3 independent groups of zebrafish, confirming the observation that zebrafish swim faster when directionally aligned.

**Data availability:** Source code 1 contains all the simulation and analysis code used to prepare this manuscript. The experimental data analyzed in this paper can be obtained at the following web address: https://drive.google.com/drive/folders/1UmzlX-yJhzQ5KX5rGry8wZgXvcz6HefD.

## Introduction

Scientists frequently look for correlations between variables to identify potentially important relationships or to support conceptual models. In disciplines with a focus on dynamics such as ecology and physiology, it is common to measure correlations between time series. Many important biological processes are nonstationary, meaning that their statistical properties (e.g., mean or variance) change systematically across time. Such processes range from the spread of an invasive species to a cell’s response to a new environmental stress.

Interpreting a correlation between a pair of nonstationary time series can be highly fraught because it is easy to obtain a seemingly impressive correlation between two time series that have no meaningful relationship [1]. For example, the sizes of any two exponentially-growing populations will be correlated over time due to a shared growth law, even if the populations lack any interactions or shared influences (e.g., bacterial cultures in separate tubes). To avoid overinterpreting spurious correlations, it helps to distinguish between the concepts of “correlation” and “dependence”. In time series research, the term “correlation” is often used procedurally [2, 3, 4]. Similarly, here we define a correlation function (*ρ*) to be any function that takes two time series and produces a statistic, although it is usually interpreted as a measure of similarity or relatedness.

In contrast to correlation, “dependence” is a hypothesis about the relationship between variables, and it has clearer scientific implications. Two events *A* and *B* are called dependent if *P* (*A, B*), the probability that they both occur, differs from *P* (*A*)*P* (*B*), the product of their individual probabilities. The formal definition of dependence extends this idea to continuous variables, where discrete probabilities are replaced by probability densities [5, 6]. Similarly, two temporal processes are dependent if the probability of observing any particular pair of trajectories differs from the product of the probabilities of the individual trajectories. The importance of dependence relationships can be attributed to Reichenbach’s common cause principle, which states that if two variables are dependent, then either they share a common causal influence, or one variable causally influences the other (possibly indirectly) [7, 8]. Thus, it is useful to first test whether an observed correlation indicates dependence before pursuing any mechanistic explanations.

There are a handful of systematic approaches to testing for dependence between nonstationary time series. However, most are fairly limited in their scope. First, when possible, a nonstationary time series can be transformed to become stationary, meaning that its statistical properties do not change systematically across time. This enables access to a wide arsenal of tests applicable to stationary data. Transformations include subtracting a trend, taking the derivative (more precisely, “differencing” between neighboring points), or choosing a stationary-seeming window of time [9, 10]. However, it is easy to see potential pathologies in each of these: Taking the derivative of an exponential curve just produces another exponential curve; subtracting a fitted linear trend from the random walk *X*(*t*) = *X*(*t* − 1) + *ϵ*(*t*) (where *ϵ*(*t*) is random noise) does not make it stationary [9]; similarly, searching for a stationary window of the same random walk process is futile since the variance of *X*(*t*) increases with each step. In a second approach, one may compare the observed correlation with correlations obtained when one time series is replaced by a synthetic replicate, where synthetic replicates are generated in a way that reflects the original nonstationary process [11]. However, these require a correct model of how the statistical properties of the process evolve over time. Lastly, there are tests within the econometrics literature that can provide evidence for dependence between random-walk-like nonstationary processes by detecting a property called cointegration, but cointegration only occurs when some linear combination of the time series is stationary [12].

Alternatively, a permutation-based approach is possible when there are multiple identically distributed and independent (iid) replicates (or trials; we use “replicates” and “trials” interchangeably). In this case, one may evaluate the significance of a within-replicate correlation by comparing it to between-replicate correlations. That is, if *X*_*i*_ and *Y*_*i*_ are time series of variables *X* and *Y* from replicate *i*, the correlation of (*X*_*i*_, *Y*_*i*_) may be compared to the correlation of (*X*_*i*_, *Y*_*j*≠*i*_). If the two variables are dependent, within-replicate correlations may differ systematically from between-replicate correlations. This approach is sometimes called “inter-subject surrogates” [13, 14].

The inter-subject surrogate approach can be used to test for dependence in each trial separately [15]. In this case, a simple nonparametric test of dependence between, for instance, *X*_1_ and *Y*_1_ can be performed by computing the correlation *ρ*(*X*_1_, *Y*_*j*_) for *j* = 1, …, *n* and writing down a *p*-value as the proportion of these correlations that are greater than or equal to *ρ*(*X*_1_, *Y*_1_) [16, 17]. For this approach, at least *n* = 20 trials are needed if one wishes to possibly obtain a *p*-value of 0.05 or below. This procedure checks for dependence in trial 1 specifically; assessing dependence in a different trial via this method would involve an analogous test.

To reduce the required number of trials below 20, a straightforward strategy is to perform a single permutation test (Fig 1A) using the data from all replicates [18, 19]. As an example with *n* = 4 trials, the test procedure begins by computing the mean within-trial correlation:

**Figure 1:**
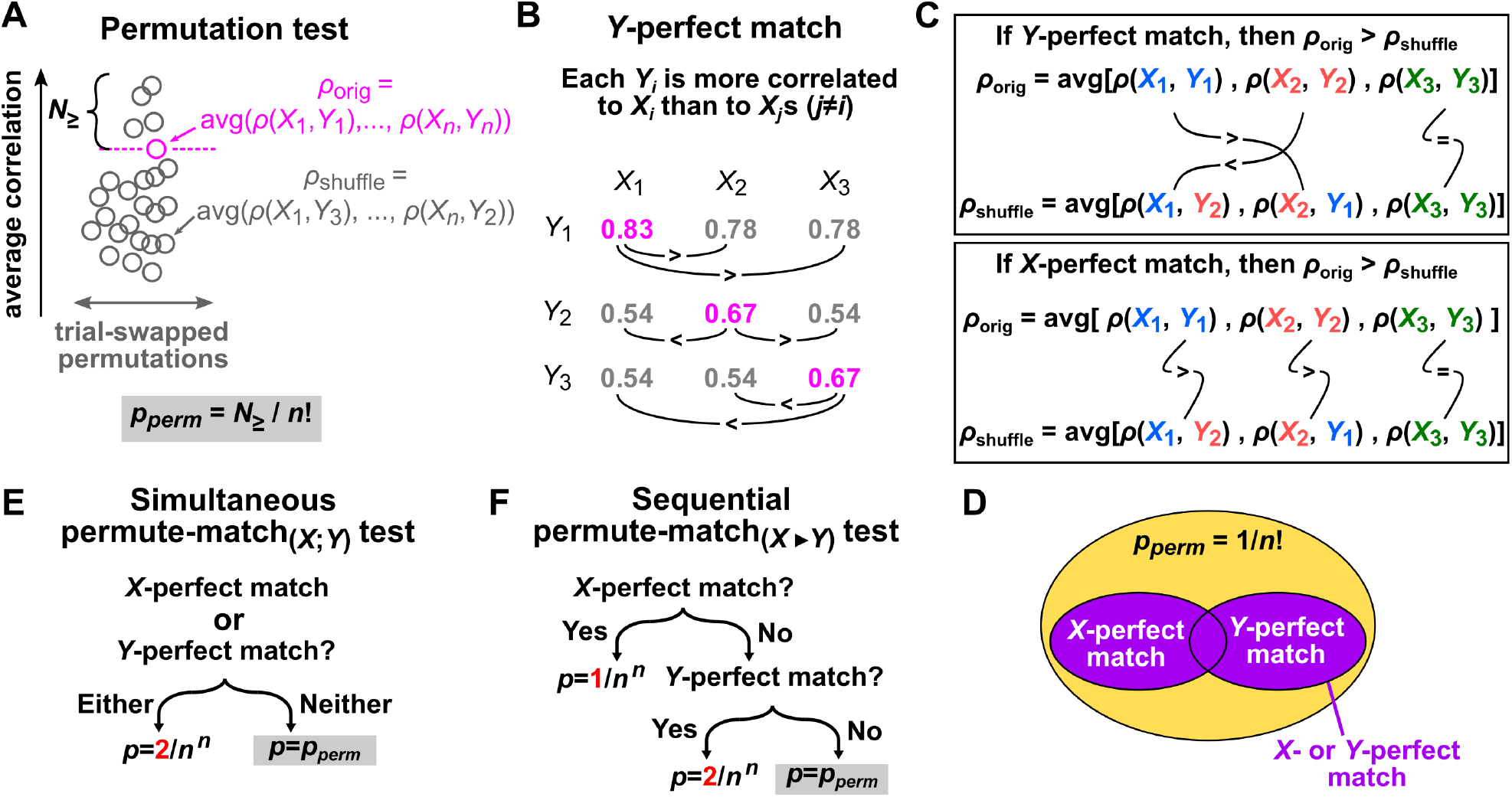
The permutation test, perfect match concept, and permute-match tests. (**A**) The permutation test. Circles indicate the average correlation without trial swapping (*ρ*_orig_; pink) or with trial swapping (*ρ*_shuffle_; grey). Here, *N*_≥_ is the number of average correlations (including *ρ*_orig_) that are equal to or larger than the original correlation *ρ*_orig_. It follows that *p*_*perm*_ = *N*_≥_/*n*! because *n*! is the number of ways the trials can be shuffled. (**B**) Illustration of a *Y*-perfect match, represented as a table of correlations wherein each diagonal entry is the greatest in its own row. Each line with a “>” or “<” symbol denotes a required ordering relationship between the two numbers at either end of the line. Note that this example has a *Y*-perfect match, but not an *X*-perfect match. For an *X*-perfect match, each diagonal entry would need to be the greatest in its own column. (**C**) If an *X*- or *Y*-perfect match occurs, then the original correlation *ρ*_orig_ is always greater than any shuffled correlation *ρ*_shuffle_. Top: If a *Y*-perfect match occurs, then each *Y*_*i*_ gives the greatest correlation when it is paired with the *X*_*i*_ from the same trial. A shuffled correlation *ρ*_shuffle_ pairs *Y*_*i*_s from some trials with *X*_*j*_s from different trials, thereby reducing the correlation. Lines with symbols “>”, “<”, and “=” denote comparisons between terms. Bottom: If an *X*-perfect match occurs, a similar argument can be applied. (D) Since a perfect match ensures that the original correlation is greater than any shuffled correlation, a perfect match also ensures that *p*_*perm*_ takes on its lowest possible value of 1/*n*!. (E) The simultaneous permute-match_(*X*;*Y*)_ test. (F) The sequential permute-match_(*X*▶*Y*)_ test; note that the permute-match_(*Y* ▶*X*)_ test is defined analogously.

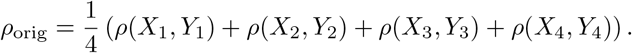

Next, as a null model, recompute the mean correlation after permuting the *Y* time series while holding the *X* time series in the original order. For instance, one such permutation might be:

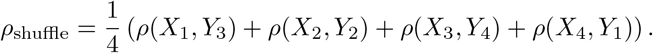

A *p*-value can be calculated as the proportion of permutations (including the original ordering) that produce a mean correlation that is as strong as, or stronger than, *ρ*_orig_ (Fig 1A). One may then detect a correlation at the significance level *α* if *p* ≤ *α*. We emphasize that these permutations are obtained by swapping trials, not by swapping time points. This test has been used to detect correlations between time series in neuroscience and psychology settings [20, 18, 21]. It has also been used to detect correlations between variables measured in brain images, which can have similar nonstationarity challenges as time series [22]. A noteworthy advantage of this approach is that it is valid (meaning that the false positive rate is guaranteed to not exceed *α*) under very mild assumptions. Namely, the test is valid if the *X*_*i*_ trials are exchangeable with one another (i.e. all permutations of the sequence *X*_1_, …, *X*_*n*_ have the same joint probability distribution), or if the *Y*_*i*_ trials are exchangeable (see Corollary 4 in Appendix 1).

Yet, even the permutation test remains considerably limited by the number of trials. The lowest *p*-value that can be obtained with *n* replicates is 1/*n*! since there are *n*! possible permutations. For example, with *n* = 3, the lowest possible *p*-value is 1/3! ≈ 0.17 and with *n* = 4, the lowest is 1/4! ≈ 0.04. These minima apply even if the average within-trial correlation is much greater than the average between-trial correlation. While replicate counts of 4 and 5 can produce *p*-values below the conventional 0.05 threshold, these *p*-values remain modest. This limitation is particularly problematic when multiple hypothesis testing is required, as the modest *p*-values may no longer be significant after applying corrections such as the Bonferroni method.

The challenge of limited replicates is a reality in many experimental settings due to cost and feasibility concerns [23, 24, 25]. Factors outside the control of investigators further restrict the number of useful replicates. For example, a study of cerebral ischemia in pigs began with 6 animals, but lost 2 due to death and experimental mishap, so complete records exist for only 4 pigs [26]. Similarly, in an animal migration study, authors affixed GPS tags to 13 juvenile cuckoos, but only 5 of them produced GPS records suitable for the study due to heterogeneous behavior among cuckoos [27]. Finally, secondary analysis of public data from earlier studies can be a resource-efficient mode of research [28], although this approach is necessarily limited to the number of replicates within the existing dataset.

In this article, we show that permute-match tests, by adding additional steps to the permutation test, can achieve *p*-values as low as 1/*n*^*n*^. For *n* = 3 or 4 respectively, the lowest possible *p*-value is then ≈ 0.04 or 0.004, compared to 1/3! ≈ 0.17 or 1/4! ≈ 0.04. The permute-match tests only require that the replicates are iid. Thus, permute-match tests allow the data analyst to detect dependence with stronger confidence when there is sufficient evidence to do so. We illustrate this approach using simulations as well as a real world example involving recordings of zebrafish swimming behavior with only 3 replicates.

## Results

### The concept of an *X*-perfect or *Y*-perfect match

Central to our approach is the concept of an *X*-perfect match or *Y*-perfect match. To motivate the ideas, suppose that a group of *n* graduate students ***X*** = (*X*_1_, …, *X*_*n*_) has been paired with a group of *n* advisors ***Y*** = (*Y*_1_, …, *Y*_*n*_) so that the *i*th student (*X*_*i*_) is paired with the *i*th advisor (*Y*_*i*_). Moreover, let *ρ*(*X*_*i*_, *Y*_*j*_) be a number that measures how well the *i*th student and the *j*th advisor get along. We say that the *i*th student is “happy” if they get along with their own advisor strictly better than they get along with any other advisor, meaning that *ρ*(*X*_*i*_, *Y*_*i*_) >*ρ*(*X*_*i*_, *Y*_*j*_) for all *j*≠ *i*. Similarly, the *i*th advisor is “happy” if *ρ*(*X*_*i*_, *Y*_*i*_) >*ρ*(*X*_*j*_, *Y*_*i*_) for all *j*≠ *i*. The arrangement of student-advisor pairs is *X*-perfect if all students are happy, and *Y*-perfect if all advisors are happy.

Analogously, if ***X*** = (*X*_1_, …, *X*_*n*_) and ***Y*** = (*Y*_1_, …, *Y*_*n*_) are collections of time series with *n* trials each, and if *ρ* is some correlation function, we say that an *X*-perfect match has occurred if (and only if) for each *i* ∈ {1, …, *n*},

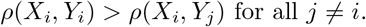

Similarly, a *Y*-perfect match has occurred if and only if for each *i*,

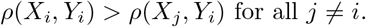

See Fig 1B for an example of a *Y*-perfect (but not *X*-perfect) match.

Throughout the article, we use boldface ***X*** or ***Y*** to indicate a collection of trials, *X*_*i*_ or *Y*_*i*_ to indicate an individual trial, and *X* or *Y* to refer to the variable in general.

A perfect match (either *X*-perfect or *Y*-perfect) provides especially strong evidence to support the hypothesis that ***X*** and ***Y*** are dependent. With a perfect match, the original correlation *ρ*_orig_ is strictly greater than any of the shuffled correlations, as illustrated in Fig 1C. Thus, a perfect match guarantees *p*_*perm*_ = 1/*n*!, the lowest *p*-value achievable with the permutation test (Fig 1D). Furthermore, letting *P* denote probability, we can prove that

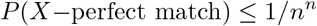

under the null hypothesis *H*_0_ wherein (1) the *Y*_*i*_ trials are iid, (2) the *X*_*i*_ trials are iid, and (3) ***X*** and ***Y*** are independent (Lemma 8 in Appendix 2). In Proposition 14 (Appendix 2), we show that this upper bound of 1/*n*^*n*^ cannot be reduced further without requiring additional assumptions.

The same bound applies to the *Y*-perfect match: under *H*_0_ we have *P* (*Y* −perfect match) ≤ 1/*n*^*n*^ (Lemma 6 in Appendix 2) and thus

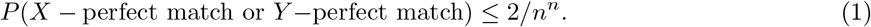

### Permute-match tests: modified permutation tests of dependence

We now describe tests for dependence between nonstationary time series based on the ideas just laid out, which we will call permute-match tests. Suppose that an experiment with *n* replicates produces time series ***X*** = (*X*_1_, …, *X*_*n*_) and ***Y*** = (*Y*_1_, …, *Y*_*n*_), and we want to test whether ***X*** and ***Y*** are dependent. For example, we may have video recordings of *n* animals, each in a separate but identically constructed enclosure; for the *i*th animal, we extract a time series of movement speed (*X*_*i*_) and a time series of the distance from a light source (*Y*_*i*_). Using these data, we may wish to investigate whether the animals under study tend to move at different speeds depending on their distance from the light source. We introduce two tests: simultaneous permute-match and sequential permute-match. As we will see, the former can be more reproducible while the latter can be more powerful.

In the simultaneous permute-match test (Fig 1E), we check for both an *X*-perfect and a *Y*-perfect match. If either the *X*- or *Y*-perfect match occurs, we write down a *p*-value to be *p* = 2/*n*^*n*^ as per equation 1. If neither *X* nor *Y* achieves a perfect match, then we fall back to the permutation test: *p* = *p*_*perm*_ = *N*_≥_/*n*! where *N*_≥_ is the number of all *n*! possible permutations (including the original ordering) that produce an average correlation that is at least as large as the original ordering (Fig 1A).

In the sequential permute-match test, we begin by choosing a variable (either *X* or *Y*) for the first perfect match test. For example, suppose we choose *X* first (the “permute-match_(*X*▶*Y*)_” test). If an *X*-perfect match is observed, then *p* = 1/*n*^*n*^ (Fig 1F). If *X*-perfect match is not observed, we then check for a *Y*-perfect match, and write *p* = 2/*n*^*n*^ if a *Y*-perfect match is observed. If neither an *X*-perfect nor *Y*-perfect match occurs, then *p* = *p*_*perm*_. A permute-match_(*Y* ▶*X*)_ test is defined analogously.

Under *H*_0_ (i.e. the *Y*_*i*_ replicates are iid, the *X*_*i*_ replicates are iid, and ***X*** and ***Y*** are independent), all the above tests are valid, meaning that for any significance level *α*, we have *P* (*p* ≤ *α*) ≤ *α*. This fact is proven as Theorems 11-13 in Appendix 2.

The permute-match tests (Fig 1E-F) are “upgrades” to the permutation test (Fig 1A) because permute-match tests always report the same or a lower *p*-value (since 1/*n*^*n*^ < 2/*n*^*n*^ ≤ 1/*n*! for all integers *n* ≥ 2; see Lemma 10 in Appendix 2). In exchange for lower *p*-values, the permute-match tests have a slightly more restrictive data requirement than the permutation test. The permutation test is valid either if the *X*_*i*_ trials are exchangeable, or if the *Y*_*i*_ trials are exchangeable (Corollary 4 in Appendix 1). The permute-match tests require that all *X*_*i*_ trials are iid and that *Y*_*i*_ trials are iid. If trials are iid, then they are necessarily exchangeable. However, exchangeable trials are not always iid. For example, the first five cards drawn from a shuffled deck are exchangeable because all possible orderings of cards are equally likely, but they are not iid because if the first card is an ace of hearts, the second card cannot be the ace of hearts. In practice, biological experimental designs typically use replicates that are both exchangeable and iid, so the more restrictive requirement of the permute-match test is often inconsequential.

The different tests – permutation test, simultaneous permute-match test, and sequential permute-match tests – are valid tests of dependence given iid replicates (Theorems 11-13 in Appendix 2). In other words, they all guarantee false positive rates at or below the significance level. Two subtle aspects of the permute-match tests are worth mentioning here. First and crucially, the permutation test and perfect match events are *nested*. That is, an *X*- or *Y*-perfect match is possible only if *p*_*perm*_ is already at its lowest possible value of 1/*n*! (Fig 1C-D). We can therefore combine the checks for perfect matches with the permutation test without inflating the false positive rate above the nominal level. This would not work if the tests were not nested, because then each test would separately contribute false positive results. The second subtlety refers specifically to the sequential permute-match tests. These tests use the lower *p*-value of 1/*n*^*n*^ if the first perfect match occurs, but use the more conservative “Bonferroni-corrected” *p*-value of 2/*n*^*n*^ if only the second match occurs (Fig 1F). This superficially resembles the practice of conducting additional tests merely because earlier tests did not result in a detection – a form of *p*-hacking that generally results in an invalid final *p*-value. However, our sequential permute-match procedure is actually valid because it combines binary events instead of continuous variables. See Appendix 3 for a detailed discussion of this point.

### Permutation and permute-match test variants may differ in statistical power

Although the permutation test, simultaneous permute-match test, and sequential permute-match test are all valid, they vary in power – the probability of detecting true dependence. Depending on the number of replicates *n* and the desired significance level *α*, there are four regimes to consider (Fig 2A).

**Figure 2:**
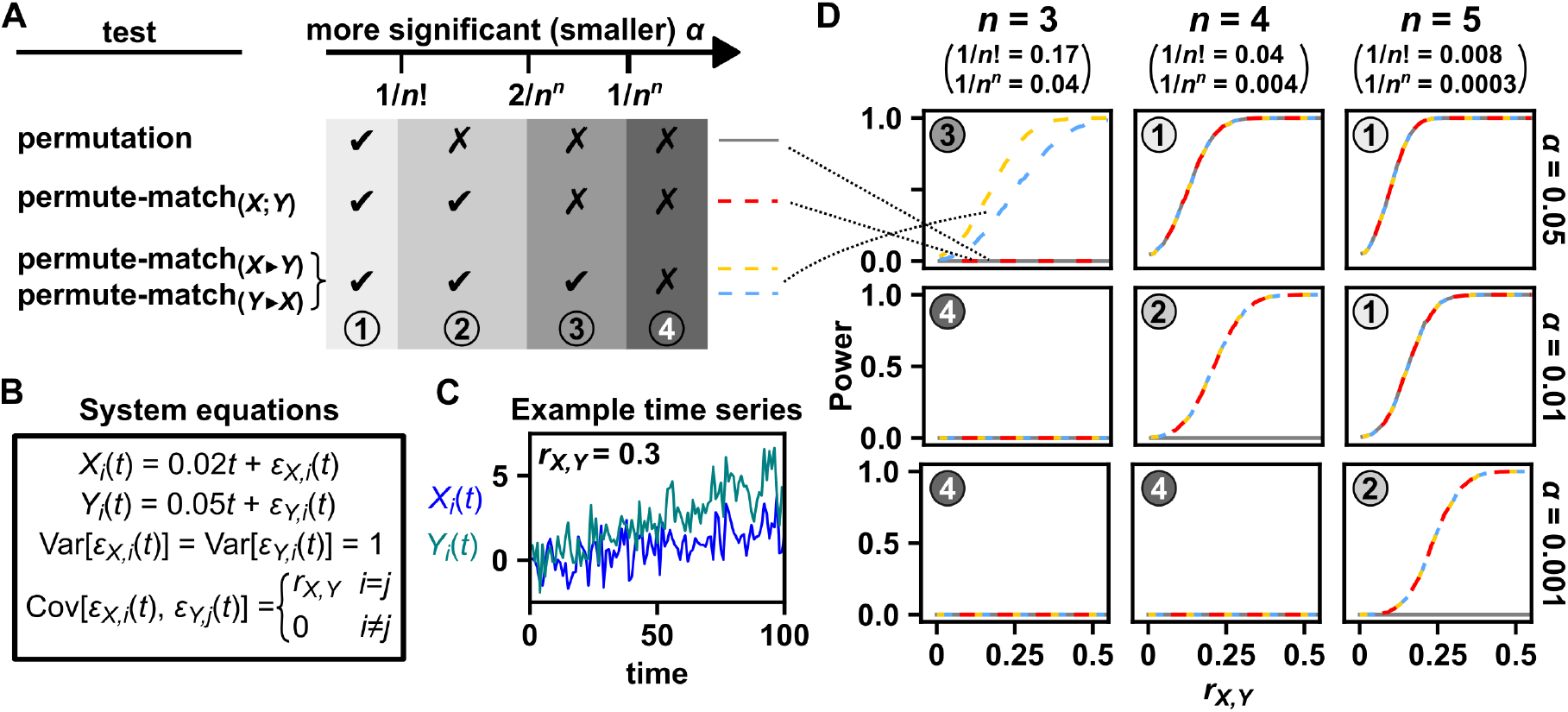
Statistical power of various permutation test variants as illustrated using a simple nonstationary system. (**A**) Summary of test power as a function of the significance level *α* and the number of replicates *n*. (**B, C**) System equations and example dynamics. The processes *X* and *Y* are given by a linear trend with additive noise on a time grid *t* = 1, 2, …, 100. The noise terms *ϵ*_*X,i*_(*t*) and *ϵ*_*Y,i*_(*t*) are drawn from a bivariate normal distribution with a mean of 0, variance of 1, and covariance of *r*_*X,Y*_. The chart shows an example pair of time series where *X* and *Y* are dependent (*r*_*X,Y*_ = 0.3). (**D**) Statistical power for the permutation test and various permute-match tests as a function of the replicate number *n*, significance level *α*, and strength of dependence *r*_*X,Y*_. Power was estimated as the proportion of simulations in which dependence was detected, calculated from 5000 simulations at each value of *r*_*X,Y*_ between *r*_*X,Y*_ = 0 and 0.54 in steps of size 0.01. At *r*_*X,Y*_ = 0, there is no dependence, so the curve at that point indicates the false positive rate rather than power. We chose the Pearson correlation coefficient as our correlation function *ρ*. See (**A**) for the color legend.

Regime 1: when *α* ≥ 1/*n*!, all tests have the same power. If the permutation test detects dependence, permute-match tests will too since they fall back to a permutation test in the “worst case” (Fig 2A, column 1). Conversely, if the permutation test did not detect dependence (i.e. *p*_*perm*_ >*α*), then it follows that *p*_*perm*_ > 1/*n*!, which implies that neither an *X*-perfect nor *Y*-perfect match occurred (Fig 1D), and so the permute-match tests cannot have detected dependence either.

Regime 2: when 1/*n*! >*α* ≥ 2/*n*^*n*^, all permute-match tests are tied for highest power (Fig 2A, column 2). In this regime, the permutation test has zero power, since its *p*-value is limited at 1/*n*!. Conversely, if either an *X*-perfect or a *Y*-perfect match occurs, then, the sequential permute-match tests as well as the simultaneous permute-match_(*X*;*Y*)_ test will all detect dependence.

Regime 3: when 2/*n*^*n*^ >*α* ≥ 1/*n*^*n*^, the simultaneous permute-match_(*X*;*Y*)_ test will have no power as the desired *p*-value is below the lowest possible *p*-value achievable by the test. Only the permute-match_(*X*▶*Y*)_ or _(*Y* ▶*X*)_ tests now have power (Fig 2A, column 3). Note that permute-match_(*X*▶*Y*)_ and permute-match_(*Y* ▶*X*)_ test may differ in power, as can be seen in the upper left panel of Fig 2D. We do not currently know how to pre-diagnose which of these two tests will be most powerful based on a given dataset, so the choice of which test to run in this case is arbitrary.

Regime 4: when *α <* 1/*n*^*n*^, no test has any power even when a perfect match occurs.

Figure 2B-D illustrates these different scenarios using a simulated nonstationary system and some common significance cutoffs. We simulated time series of two variables (*X* and *Y*) from a small number of replicates (between 3 and 5). Here, the time series were produced from a pair of linear time trends with coupled additive noise (*r*_*X,Y*_ determines the strength of coupling; Fig 2B). We chose the Pearson correlation coefficient as our correlation function *ρ*. The various sub-panels of Fig 2D illustrate the different aforementioned regimes.

The above example was chosen for ease of presentation: in fact, the example is so simple that it may be reasonably treated by just fitting and removing the linear trend. In Appendix 4, we show the overall higher statistical power of permute-match tests over the permutation test in a more complex example where the data-generating process is nonstationary (a logistic map with time-varying parameters) and where a nonlinear correlation function (cross-map skill) is preferable to (linear) Pearson correlation.

### The permute-match test’s false positive rate depends on the process tested

The permute-match test is conservative – meaning that its false positive rate can fall below the significance level – because both the permutation test and perfect match test are conservative. As discussed elsewhere [29], the permutation test is conservative when *α* is not one of its possible *p*-values and when ties may occur between the original correlation and shuffled correlations. Checking for a perfect match is similarly conservative. The actual probability of a false-alarm perfect match event can vary depending on the process being tested.

To see this consider the permute-match_(*Y* ▶*X*)_ test in the setting where *n* = 3, where *α* = 0.05, and where *r*_*X,Y*_ = 0 (independent *X* and *Y*). Note that in this case, obtaining a *Y*-perfect match (and thus *p* = 1/*n*^*n*^) is necessary and sufficient to detect dependence since *α* is too low for detection by either the permutation test or the *p* = 2/*n*^*n*^ leg of the permute-match test. In the linear system of Fig 2, we observed among 5000 simulations a detection rate of 0.0148, significantly below the upper bound of 1/3^3^ (Figure 2 – Source data 1; one-tailed exact binomial test, *p <* 10^*−*20^). Conversely, in the nonlinear system of Fig S2, this same event (*Y*-perfect match under *n* = 3 and *r*_*X,Y*_ = 0) occurs with a detection rate of 0.0328, not significantly below the upper bound of 1/3^3^ (Figure S2 - Source data 1; one-tailed exact binomial test, *p* = 0.059). Thus, depending on the underlying process studied, the actual chance of a perfect match happening under independence may be near or significantly below the theoretical upper bound.

### An example from animal behavior science

Living in groups is a common experience among animals. For example, at least half of fish species are thought to form groups at some stage of life [30, 31]. The properties of these groups can impart a variety of important fitness effects [32]. In groups of fish, coordinated swimming may facilitate foraging (e.g., by enabling fish to “communicate” the location of food to others) and predator escape (e.g., by confusing predators) [33, 31]. One study [31] observed, in videos of small groups of zebrafish (8 fish per group), that the fish swam faster when in a high-polarization state (i.e. directionally aligned) than when in a low-polarization state. A correlation between polarization and speed might indicate statistical dependence between the two variables, or it might just be a consequence of temporal trends (e.g., polarization and swimming speed both happen to decrease with time).

Using a publicly available dataset [34] containing fish trajectories from 3 replicate 10-minute videos of small groups of juvenile *Danio rerio* zebrafish (10 fish per group), we performed a permute-match test of dependence between polarization and fish swimming speed. As we will see, although swimming speed and the extent of polarization are potentially nonstationary, the permute-match test nevertheless detects a statistical dependence between them.

Directional polarization is quantified as the mean resultant length *C*_*t*_ of fish velocity [35]. We represent the swimming direction of each fish by a unit vector, and then define *C*_*t*_ as the length of the average unit vector (Fig. 4; Eq. 2). The mean resultant length is bounded between 0 (e.g., two opposite vectors) and 1 (e.g., two parallel vectors). To quantify speed, we took the “average individual speed”, which is a term we use to denote an average across individuals, not across time. More precisely, the average individual speed of a 10-fish group at time *t* is (*s*_1,*t*_ + *s*_2,*t*_ + · · · + *s*_10,*t*_)/10 where *s*_*i,t*_ is the speed of fish *i* at time *t*. Note that average individual speed is distinct from the speed of the group center, which has also been studied in *Danio* fish [32].

Neither the average individual speed nor the mean resultant length is obviously stationary (Fig 3A). By visual inspection, the average individual speed seems to decrease across time, and all time series are deemed nonstationary by a Kwiatkowski-Phillips-Schmidt-Shin test (*p <* 0.01), and so we may be on shaky ground to use methods that require stationarity. Additionally, since only 3 trials are available, the permutation test and the simultaneous permute-match test cannot detect dependence at the 0.05 level, so we perform a sequential permute-match test.

**Figure 3:**
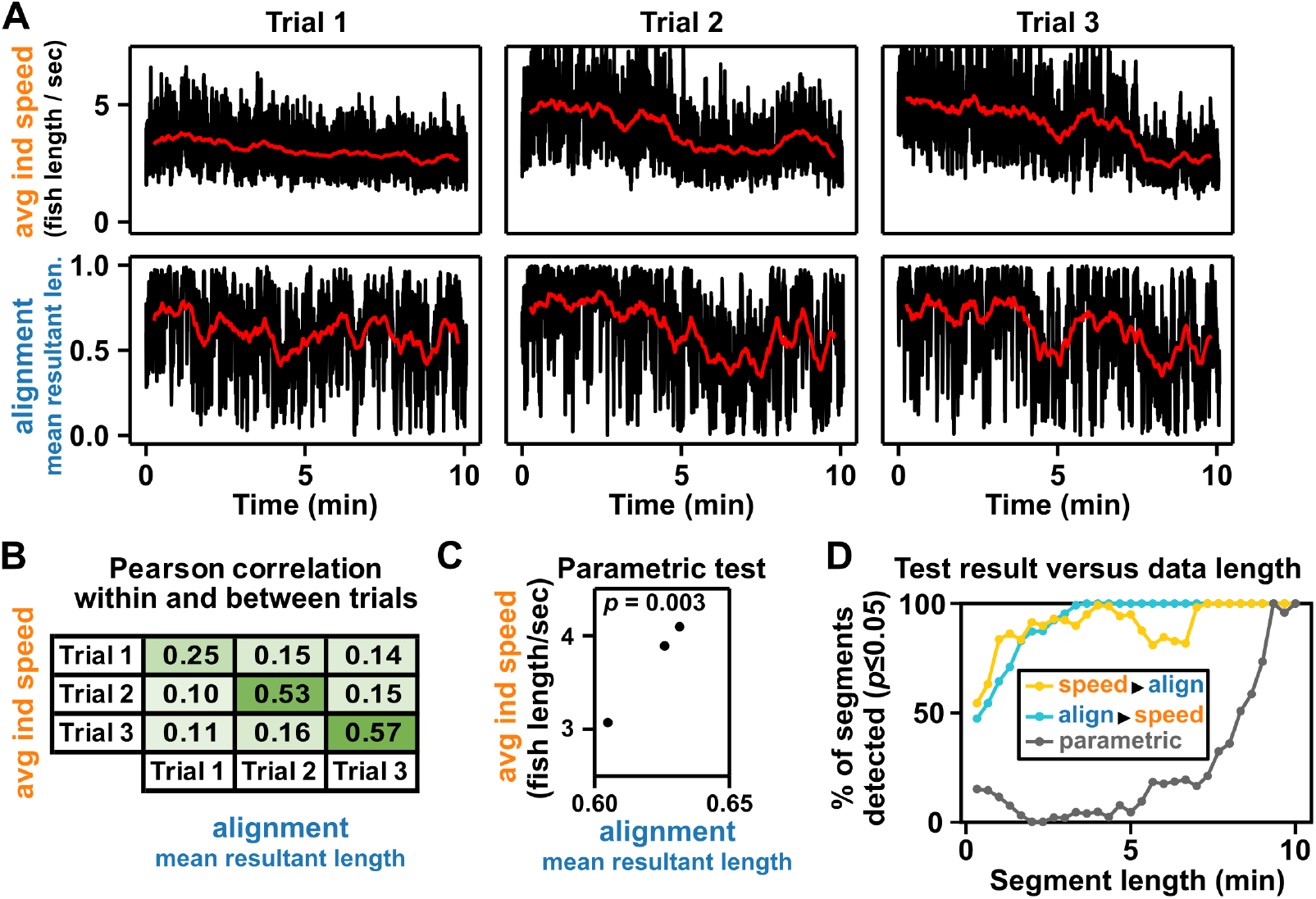
A sequential permute-match test detects dependence between average individual speed and alignment (as measured by mean resultant length) in small groups of zebrafish. (**A**) Time series of average individual speed and mean resultant length for the three replicate videos. Black curves show original time series, and red curves show a 30-second moving average. Average individual speed appears to decrease with time in all trials. All 12 time series (2 variables × 3 trials × 2 smoothing conditions) were deemed non-stationary by a Kwiatkowski-Phillips-Schmidt-Shin (KPSS; [37]) test (*p <* 0.01). Note that the KPSS test seeks to reject a stationary null hypothesis in contrast to other common tests, whose null hypotheses are nonstationary. (**B**) Pearson correlation between alignment and average individual speed, both within trials and between trials, using all frames up to the length of the shortest speed and alignment time series (19308 frames; 10.06 minutes). Table entries are shaded by correlation. We observe a speed-perfect match (and an alignment-perfect match), and thus detect dependence with *p* = 1/3^3^ ≈ 0.04. (**C**) A parametric test also detects dependence. As a parametric alternative, for each trial we averaged values of speed and mean resultant length over the complete time series (visualized in the scatter plot shown, where each point is one trial). The sample Pearson correlation of the time-averaged variables is 0.99996, and a one-tailed test of significance gives *p* ≈ 0.003. (**D**) Permute-match tests detected a significant correlation between speed and alignment more consistently than the parametric test. For a grid of lengths between 20 and 600 seconds we sampled 500 random segments of each length, each drawn from the first 600 seconds, and determined for each segment whether the parametric test and/or the two possible permute-match tests detected a significant (*p* ≤ 0.05) correlation. In the edge case of the maximum 600-second length, all 500 “random” segments were identical. We used the Pearson correlation as the correlation function *ρ* in the permute-match tests, and a one-tailed parametric test, reflecting the alternative hypothesis of correlation being positive (not merely nonzero).

**Figure 4:**
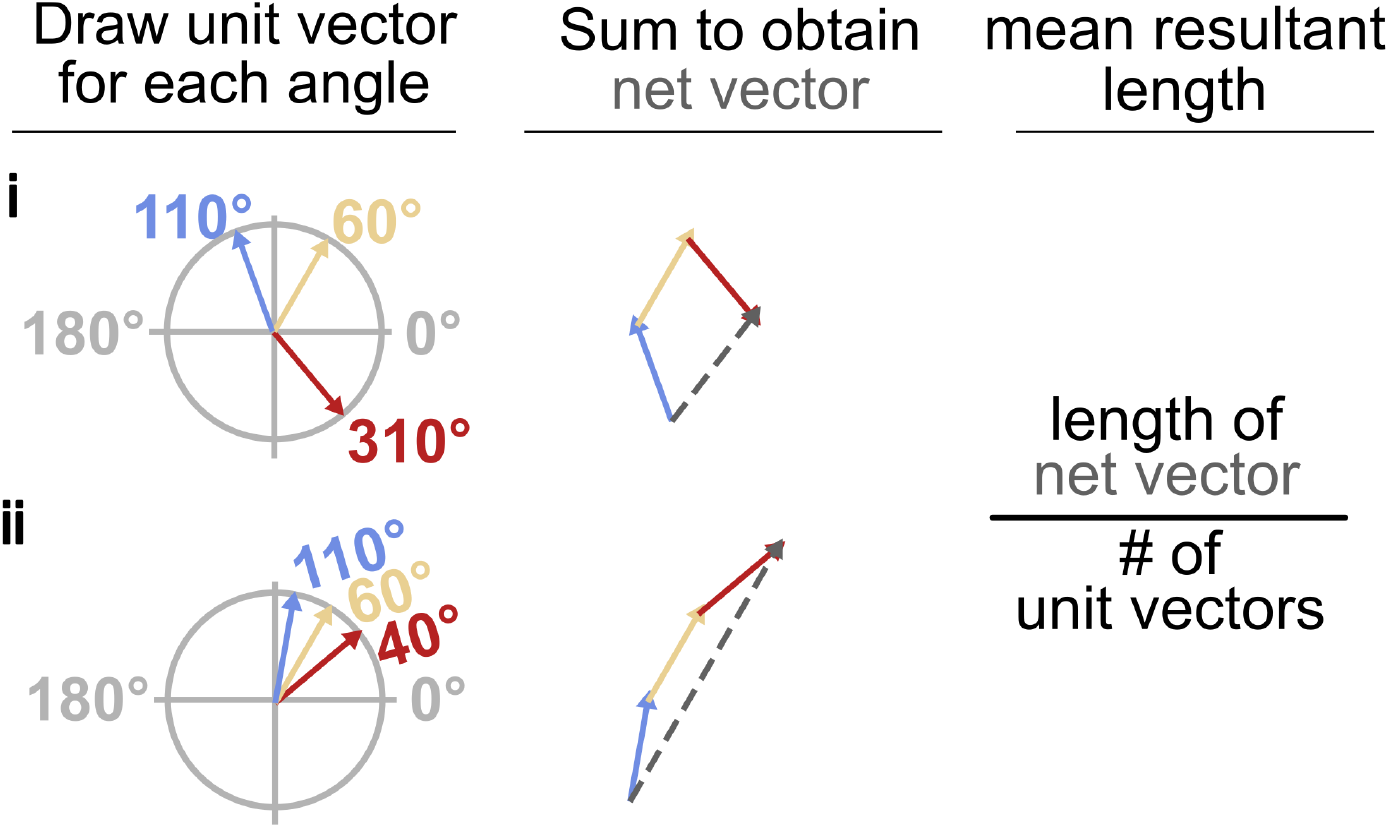
The mean resultant length is a measure of alignment among angles. Consider three unit vectors colored blue, yellow, and red. The length of net vector (dashed) divided by the number of vectors is defined as the mean resultant length. Compared to less-aligned angles (i), strongly aligned angles (ii) produce a longer net vector, and thus a greater mean resultant length.

We arbitrarily started with average individual speed and identified a “speed”-perfect match, indicating a significant correlation between average individual speed and mean resultant length (Fig 3B; *p* = 1/3^3^ ≈ 0.04). Significance does not depend on the choice of which match is tested, since both a “speed”-perfect and a “mean resultant length”-perfect match are obtained. Note that although trials are independent of each other, all between-trial correlations from the full dataset are positive (off-diagonal elements in Fig 3B), illustrating the danger in applying naive data analysis methods to data that are autocorrelated or even nonstationary.

To try a parametric alternative, for each trial we time-averaged the average individual speed and mean resultant length over the full 10 minutes (Fig 3C; where each point is one trial). The sample Pearson correlation of the time-averaged variables is 0.99996, and a one-tailed test of significance (using the function stats.pearsonr from the Python package Scipy [36]) also detects dependence (*p* ≈ 0.003), consistent with the permute-match tests. This test has the null hypothesis that the two variables shown in Fig 3C (time-averaged average individual speed and time-averaged mean resultant length) are independent and Gaussian. We view this test as comparable to the permute-match tests because although it requires a distributional assumption, it can be valid even when the time series are nonstationary.

Since data are limited in many scientific applications, we explored whether the results of the sequential permute-match tests and the parametric Pearson correlation test were consistent across different time series lengths. We truncated time series to various lengths (from 20 seconds to 10 minutes), and checked whether each test reported a significant (*p* ≤ 0.05) correlation (Fig 3D). Both sequential permute-match tests reported a significant correlation at all lengths tested, whereas the parametric test was inconsistent, finding a significant correlation for about one third of lengths.

## Discussion

Outside the context of time series, permutation tests have been celebrated because they require only minimal assumptions [38]. To satisfy the requirements of permutation tests, it is sufficient (see Corollary 4 in Appendix 1) to collect data from exchangeable replicates. Data in each trial can be autocorrelated (e.g., temporally or spatially) and nonstationary. No distributional assumptions are required, and the correlation function *ρ* can be completely arbitrary. For instance, *ρ* could include a lag to detect delayed dependence, or even evaluate the correlation strength at several lags and report the strongest among them [39]. Moreover, the *ρ* can be asymmetric so that *ρ*(*X, Y*)≠ *ρ*(*Y, X*) as occurs when correlations are based on techniques that use one time series to predict values of another (e.g., Appendix 4). Such freedom from assumptions stands in contrast to the situation in the analysis of single-replicate time series, where the practitioner may be forced to make assumptions that are difficult to verify. For example, statistical tests of correlation between two time series often require stationary data at a minimum [40]. Yet, it is difficult to verify that a single time series is stationary because stationarity is a property of an entire ensemble of time series [40], so checking for stationarity in a single time series is similar in spirit to checking whether a single data point is drawn from a Gaussian distribution.

However, applying a trial-swapping permutation test to check for significant correlations requires that a sufficient number of trials be available to permute amongst one another. When only small numbers of trials are available, the permutation test is severely limited by mathematics. The minimum *p*-value that can possibly be achieved by a permutation test is 1/*n*!, which will provide only modest evidence against a null hypothesis when *n* = 4 and is essentially useless when *n* = 3, regardless of how strong the true dependence may be. This limitation is particularly severe for researchers testing several related hypotheses, as this ideally requires a test that can produce *p*-values sufficiently low to maintain significance even after a multiple-testing correction.

Here we have described modified permutation tests – the sequential and simultaneous permute-match tests – that have access to lower *p*-values. In the sequential permute-match test, if the first variable does not display a perfect match, this test falls back to the simultaneous permute-match test. Thus, the sequential permute-match test is more powerful than the simultaneous test. However, the sequential test requires the arbitrary choice of checking for an *X*- or *Y*-perfect match first, making it potentially less reproducible between studies than the simultaneous test, which does not involve an arbitrary choice. If neither an *X*-nor *Y*-perfect match is obtained, both tests default to a permutation test. This step is valid because the tests are nested in the sense that a perfect match can only occur when the permutation *p*-value takes on its lowest possible value (Lemma 9 in Appendix 2).

The test appears computationally tractable for relevant sample sizes: Our implementation of the permute-match_(*X*▶*Y*)_ procedure completed a single test of dependence in the setting of Fig 2 with an average runtime of 3 seconds when *n* = 10 on a 2023 14-inch MacBook Pro with an M2 Pro processor and 16 GB RAM (see Source data 1). For *n >* 10, a standard permutation test already can report a *p*-value below 3 × 10^*−*8^, so the perfect match test is likely unnecessary for typical applications.

It seems likely that the tests could be further modified to report an even lower *p*-value when an *X*- and *Y*-perfect match occur simultaneously, as in Fig 3B. However, we have not investigated further and this problem is left for future efforts.

Although here we focused on time series, the permute-match test can in principle be applied to other scenarios where iid replicates are available, but data within each replicate are dependent. One example is spatial data in brain imaging, where the permutation test has been performed using multiple scanned brains [22]. Beyond spatial processes, dependent data also appear in nucleotide sequences and natural language text. Overall, advances in methods that take advantage of multiple replicates may facilitate statistical testing without requiring assumptions that are difficult to verify.

## Methods

### Data preprocessing

We obtained fish trajectory data [34] from the web address at https://drive.google.com/drive/folders/1UmzlX-yJhzQ5KX5rGry8wZgXvcz6HefD. For the first trial we used the file at the path “10/1/trajectories_wo_gaps.npy” (where “10” indicates the number of fish and “1” indicates the first trial). For the second and third trials, the file path was the same except it began with “10/2/” and “10/3/” respectively. These files contain sequences of fish positions indexed by video frame, as well as constants such as the frame rate (32 frames per second) and approximate fish body length.

We estimated the velocity of each fish as

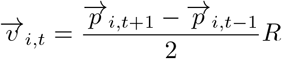

where 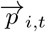 is a 2-dimensional column vector specifying the position of the *i*th fish in units of body length at video frame *t*, 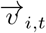 is a 2-dimensional column vector specifying the fish velocity in units of body length per second at video frame *t*, and *R* is the frame rate (32 frames per second). The 3 trials contained similar but slightly different numbers of frames. We truncated the 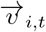 time series to the shortest among them (19308 frames ≈ 10.06 minutes) so they could be compared before downstream analyses.

A group’s average individual speed at each time was then given by

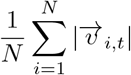

where *N* is the number of fish. The mean resultant length *C*_*t*_ of a group at each time was given by the formula

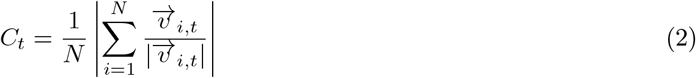

Velocity vectors 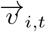 covering missing tracking data (fewer than 0.02% of such vectors in any trial) were excluded from calculations of average individual speed and mean resultant length. When calculating mean resultant length, fish with speed of exactly zero (whose direction of motion is undefined) were also excluded. Instances of zero speed were rare, occurring in at most 1 fish-frame in any trial. Average individual speed and mean resultant length were determined from the remaining fish (always at least 8 of 10), so no time point was dropped from any trial.

### Statistical analysis

The permute-match test is described in Fig 1 and its validity is proved in Appendix 2. For the KPSS test, we used the function “statsmodels.tsa.stattools.kpss” in the Python package Statsmodels (ver. 0.10.1) [41]. We set the “regression” argument to “c” (which tests the null hypothesis of stationarity rather than trend-stationarity), and we set the “lags” argument to “auto” (which means that the number of lags used for testing stationarity is automatically chosen by a data-dependent method).

### Artificial intelligence

ChatGPT 5.2 was used while constructing the proof of Lemma 15, particularly in the application of the “taking out what is known” property. Claude Opus 4.8 was used to review code, mathematical arguments, and article text for errors and unclear language. Claude Opus was also used to edit some parts of the manuscript for brevity. Corrections suggested by Claude Opus were reviewed by the authors and incorporated where appropriate.

## Acknowledgements

We thank Yunyi Zhang (CUHK-Shenzen) as well as members of the Shou Lab and two anonymous reviewers for valuable critiques. The University College London Computer Science High Performance Computational Cluster (https://hpc.cs.ucl.ac.uk/) provided computational resources and support that contributed to the completion of this research.

## Additional files

**Figure 2 - Source data 1:** Numerical values shown in Figure 2D

**Figure 3 - Source data 1:** Numerical values shown in Figure 3D

**Figure S2 - Source data 1:** Numerical values shown in Figure S2D

**Figure S3 - Source data 1:** Numerical values shown in Figure S3

**Source data 1:** Permute-match test run time as a function of the number of replicates

**Source code 1:** Python scripts used to produce the results shown in Figures 2, 3, S2, and S3 as well as Source data 1.

## Appendix

In Appendix 1, we review the basic permutation test of independence and a proof of its validity. This is because, although these results are known to the statistical community, they are often presented in an abstract form whose connection to independence testing may not be obvious to the nonspecialist, or parts of the argument are simply left as an exercise to the reader. In Appendix 2, we provide the theoretical justification for the permute-match test, which is the main topic of this work. Appendix 3 expands on a brief claim made in the main text about *p*-values of sequential statistical tests, and Appendix 4 contains supplementary figures.

### 1 Justification of the permutation test of independence

In this section we provide a proof that the permutation test of independence is valid. No major novelty is claimed in this subsection. For instance, the results of Lemma 3 can be mostly reconstructed by piecing together various theorems, examples, and homework problems from section 15.2.1 of Lehmann and Romano [42]. Nevertheless, we present the complete argument here as a courtesy to the reader. Related proofs or proof sketches of various kinds of permutation tests can be found in [19, 29].

The following lemma describes a general rank-based method to test whether the distribution of a random variable changes upon applying a set of transformations. This lemma and proof are based on Theorem 15.2.1 in Lehmann and Romano [42]. We begin with this result because it comes first in the chain of logical steps that justify the permutation test, but it is somewhat abstract, and so the reader who prefers to start out with concrete concepts may wish to skip to Def 2 and return to this lemma when its necessity becomes apparent later. As a notational clarification, note that for the transformation *h*_*i*_, to denote “*h*_*i*_ applied to *Z*” we write *h*_*i*_*Z* rather than the more traditional *h*_*i*_(*Z*).

#### Lemma 1

Let *Z* ∈ Z be a random variable. Let *H* = {*h*_1_, …, *h*_*m*_} be a set of functions from Z to Z such that:

1. the probability distributions of *Z* and *h*_*i*_*Z* are the same for all functions *h*_*i*_ in *H*, and
2. the unordered sets {*h*_1_*Z*, …, *h*_*m*_*Z*} and {*h*_1_*h*_*i*_*Z*, …, *h*_*m*_*h*_*i*_*Z*} are equal for all *h*_*i*_ in *H*.

Let the statistic *T* be a function from *Ƶ* to ℝ. Given some *z* ∈ *Ƶ*, let the ordered values of {*T* (*h*_1_*z*), …, *T* (*h*_*m*_*z*)} be

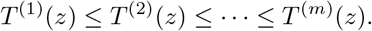

Choose some significance level *α* ∈ [0, 1] and let the rank *k* be

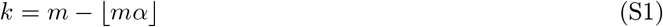

where ⌊*mα*⌋ is the largest integer less than or equal to *mα*. Then,

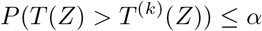

where *P* (·) denotes the probability of an event.

#### Remark

Before proceeding to the proof, let us walk through the lemma, using a concrete example to help with the most abstract aspects. Suppose that random variable *Z* is the result of a fair coin flip. That is, *Z* = 0 with 50% chance and *Z* = 1 with 50% chance. Then *Ƶ* (the set of valid *Z* values) is the set 0, 1, and *z* can take on the value of 0 or 1. Next the lemma introduces *H*, which is a set of functions that need to satisfy certain requirements with respect to *Z*. Let’s choose *H* = *h*_1_, *h*_2_ where *h*_1_ “turns the coin over” and *h*_2_ leaves the coin as it is. That is, *h*_1_*z* = 1 − *z* and *h*_2_*z* = *z*. To check that *H* satisfies requirement (1), note that flipping the fair coin over does not change the probability distribution of its outcome, and neither does leaving it alone. To see that *H* satisfies requirement (2), we need to check that

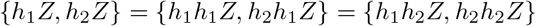

If *Z* = 0, then the above expression becomes

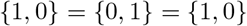

and if *Z* = 1, then it is

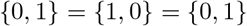

and since these are unordered sets, they are all equal, so we have verified that *H* satisfies requirement (2). Next, we need a function *T*. For example, choose *T* (*z*) = *z* + 10. Thus, for *z* = 0 we have {*T* (*h*_1_*z*), *T* (*h*_2_*z*)} = {11, 10}. The lemma then introduces a ranking notation where *T* ^(*i*)^(*Z*) is the *i*th smallest term in the sequence. This notation is useful as we will be dealing with the set {*T* (*h*_1_*Z*), …, *T* (*h*_*m*_*Z*)} for potentially large values of *m*. Returning to our example, if *z* = 0, then *T* ^(1)^(0) = 10 and *T* ^(2)^(0) = 11. Similarly for *z* = 1, *T* ^(1)^(1) = 10 and *T* ^(2)^(1) = 11. Also, in the statement of the lemma, *T* ^(*i*)^ is defined as a function of *z*, not *Z*. This is to emphasize that the ordering depends on the particular outcome of the random variable and is not fixed. Finally, the lemma makes a claim about the probability that *T* (*Z*) exceeds *T* ^(*k*)^(*Z*), which is the *k*th lowest statistic in the set. In the example, if *α* = 0.05, then *k* = 2 − ⌊ 0.1 ⌋ = 2. *T* ^(2)^(*Z*) is always going to be the second smallest, i.e. the largest, element of *T* (*h*_1_*z*), *T* (*h*_2_*z*), which is 11. Of course, the probability of *T* (*Z*) being *even greater* than the largest number is 0, and since 0 is less than *α* = 0.05 we have verified the lemma in this case. As another example, suppose *α* = 0.5. Then *k* = 2 − ⌊1⌋ = 1. *T* ^(*k*)^(*Z*) is then *T* ^(1)^(*Z*), the smallest element of {*T* (*h*_1_*z*), *T* (*h*_2_*z*)}. The probability of *T* (*Z*) being greater than the smallest number is 0.5 ≤ *α* = 0.5, which verifies the lemma for *α* = 0.5.

**Proof of Lemma 1:** Let *I*(*A*) be the indicator function for the event *A* (meaning that *I*(*A*) = 1 if *A* occurs and *I*(*A*) = 0 otherwise).

First suppose that *T* ^(*k*)^(*Z*) is not tied with any other *T* ^(*i*)^(*Z*). Then Eq. S1 implies that there are ⌊ *mα*⌋ values of *T* ^(*i*)^(*Z*) that are greater than *T* ^(*k*)^(*Z*). That is,

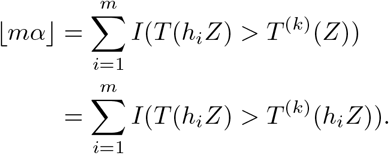

To arrive at the second equality, we used the fact that *T* ^(*k*)^(*Z*) = *T* ^(*k*)^(*h*_*i*_*Z*), which follows from the second requirement of the lemma (i.e. {*h*_1_*Z*, …, *h*_*m*_*Z*} = {*h*_1_*h*_*i*_*Z*, …, *h*_*m*_*h*_*i*_*Z*}).

Alternatively, if there is possibly a tie for *T* ^(*k*)^(*Z*) (meaning that there may be some *j ≠ k* such that *T* ^(*j*)^(*Z*) = *T* ^(*k*)^(*Z*)), then we have a similar formula, but with an inequality instead of an equality:

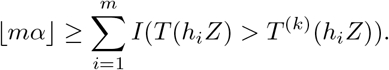

This version with the inequality is of course general since it is true under both cases. Using this formula,

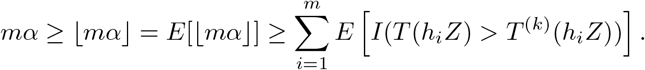

Since *Z* and *h*_*i*_*Z* have the same distribution, we may remove the *h*_*i*_ in the above formula, and we have

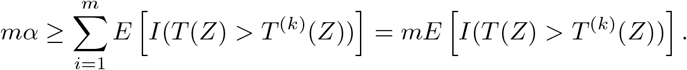

Dividing through by *m* gives

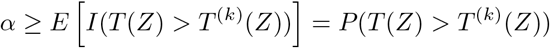

as required.

Next we give a definition of the permutation test. The definition here is somewhat more general than in the main text. As with *h*_*i*_, we will denote “*g*_*i*_ applied to *Z*” by writing *g*_*i*_*Z*.

#### Definition 2

*The permutation test of independence*

Let ***X*** = (*X*_1_, …, *X*_*n*_) and ***Y*** = (*Y*_1_, …, *Y*_*n*_) be sequences of random variables. Let *G* = {*g*_1_, …, *g*_*n*!_} be the complete set of functions that permute a sequence of length *n*. For example, if ***Y*** is (2, 7, 4), then *g*_*i*_***Y*** might be (2, 4, 7). Define the statistic *T* (***X, Y***) to be a function that returns a real number, typically interpreted as the overall correlation strength. Repeatedly compute the statistic after permuting the elements of ***Y***. That is, compute

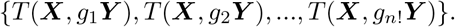

Let *N*_≥_ be the number of permuted statistic values that are at least as large as the original statistic value. That is,

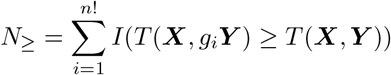

where *I*(*A*) is the indicator function for the event *A* (meaning that *I*(*A*) = 1 if *A* occurs and *I*(*A*) = 0 otherwise). Then the *p*-value of the test is given by

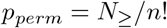

We now prove the validity of the permutation test.

#### Lemma 3

*The permutation test of independence is valid when* ***Y*** *is exchangeable*.

Let ***X*** = (*X*_1_, …, *X*_*n*_) and ***Y*** = (*Y*_1_, …, *Y*_*n*_) be sequences of random variables such that ***Y*** is exchangeable and where ***X*** is independent of ***Y***. Perform the permutation test of independence (Def. 2) to obtain *p*_*perm*_. Then, *P* (*p*_*perm*_ ≤ *α*) ≤ *α* for any *α* ∈ [0, 1].

**Proof:** Here we use the symbols *g*_*i*_ and *T* with their meanings from Def. 2.

We first set up the problem in a way that allows us to apply Lemma 1. To construct the random variable *Z* and transformation set *H* for Lemma 1, we choose *Z* = (***X, Y***) and *h*_*i*_*Z* = (***X***, *g*_*i*_***Y***). Lemma 1 has two requirements. First, *Z* and *h*_*i*_*Z* must have the same distribution. This requirement is satisfied: Since ***Y*** is exchangeable and ***X*** is independent of ***Y***, it follows that (***X, Y***) has the same distribution as (***X***, *g*_*i*_***Y***). The second requirement is that {*h*_1_*Z*, …, *h*_*m*_*Z*} = {*h*_1_*h*_*i*_*Z*, …, *h*_*m*_*h*_*i*_*Z*}. This requirement is also satisfied: The set of permutations of ***Y*** is the same as the set of permutations of *g*_*i*_***Y***. Since both requirements are satisfied, we may apply Lemma 1.

We now prove the result directly. Below, *A* ↔ *B* means “*A* if and only if *B*”.

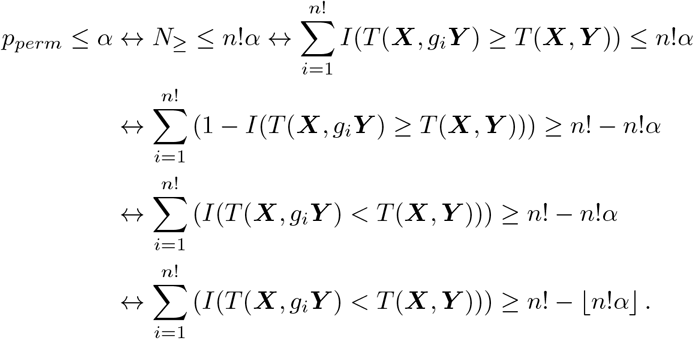

As in Eq. S1, let us define *k* = *m* − ⌊ *mα* ⌋, where *n*! = *m*. Then, the final line above says “the number of permuted ***Y*** s that produce a smaller *T* than the original ***Y*** is greater than or equal to *k*.” Said another way, “when we rank the *T* s from least to greatest, the *T* from the original ***Y*** will have a higher rank than *k*”. This in turn is equivalent to *T* (***X, Y***) >*T* ^(*k*)^(***X, Y***). Thus, we may directly apply Lemma 1, thereby obtaining

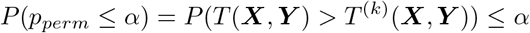

as required.

Lemma 3 is “asymmetric” in that it requires ***Y*** to be exchangeable and says nothing about ***X***. It is possible to write down a variant of this lemma where either an exchangeable ***Y*** or an exchangeable ***X*** is sufficient, but this requires an extra condition on *T*. Specifically, it is required that applying the same permutation to both ***X*** and ***Y*** must leave the value of *T* (***X, Y***) unchanged. That is, *T* (***X, Y***) = *T* (*g*_*i*_***X***, *g*_*i*_***Y***) for any permutation function *g*_*i*_. Importantly, the form of *T* that is used for the permute-match test (i.e. 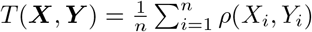) satisfies this requirement because in this case, *T* (*g*_*i*_***X***, *g*_*i*_***Y***) simply rearranges the order of the terms in the summation, which clearly has no effect on the value of *T*. We establish this formally in the corollary below.

#### Corollary 4

*If T is “nice”, then the permutation test is valid when either* ***X*** *or* ***Y*** *is exchangeable*.

Let ***X*** = (*X*_1_, …, *X*_*n*_) and ***Y*** = (*Y*_1_, …, *Y*_*n*_) be sequences of random variables such that at least one of (***X, Y***) is exchangeable and where ***X*** is independent of ***Y***. Perform the permutation test of independence (Def. 2) using a correlation function with the “nice” property that

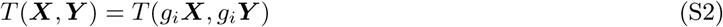

for any permutation function *g*_*i*_. Then, *P* (*p*_*perm*_ ≤ *α*) ≤ *α* for any *α* ∈ [0, 1].

**Proof:** Lemma 3 already covers the case where ***Y*** is exchangeable. We now consider the case where ***X*** is exchangeable, but not ***Y***.

We proceed by showing that the permutation test where ***Y*** is permuted is equivalent to the permutation test where ***X*** is permuted. Notice that to show this equivalence it is sufficient to establish that

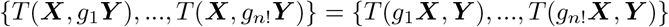

where the left and right sides both denote unordered sets. It will be useful to refer to the inverse permutation 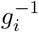, which is the permutation that “undoes” *g*_*i*_. More precisely, 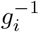 is defined by the relation 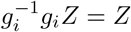. Using Eq. S2, we have:

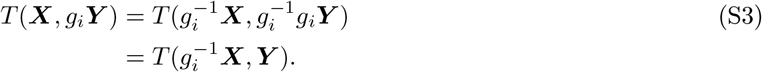

Note that the inverse permutation 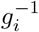 is guaranteed to exist because any permutation *g*_*i*_ can be “undone” by some other permutation 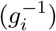. In the edge case where *g*_*i*_ is the trivial identity permutation, 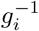 is also the identity permutation. Additionally, the set of permutation functions is the same as the set of inverse permutation functions, meaning that

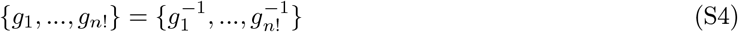

because each permutation is an inverse permutation (because *g*_*i*_ inverts 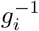), and each inverse permutation is a permutation. It follows that

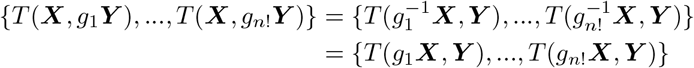

where the first equality follows from Eq. S3 and the second equality follows from Eq. S4. This shows that the permutation test where ***Y*** is permuted is equivalent to the permutation test where ***X*** is permuted.

From Def. 2 and Lemma 3 it is clear that a permutation test where ***X*** is permuted instead of ***Y*** would be valid (i.e. *P* (*p*_*perm*_ ≤ *α*) ≤ *α*) if ***X*** is exchangeable. Since we have just now shown that the present procedure is equivalent to a permutation test where ***X*** is permuted instead of ***Y***, it follows that this procedure is valid when ***X*** is exchangeable, which completes the proof.

### 2 Validity of the permute-match test

We now justify the validity of the permute-match test, which is the primary contribution of the present work. Recall from the main text that the null hypothesis of this test is that *X*_1_, …, *X*_*n*_ are iid, *Y*_1_, …, *Y*_*n*_ are iid and ***Y*** is independent of ***X***.

#### Definition 5

*The perfect match events (M*_*X*_ *and M*_*Y*_ )

Let ***X*** = (*X*_1_, …, *X*_*n*_) and ***Y*** = (*Y*_1_, …, *Y*_*n*_) be sequences of random variables. Let *ρ* (the “correlation function”) be a function that maps a pair (*X*_*i*_, *Y*_*j*_) to a real number. We say that the event *M*_*Y*_ (also called a “*Y*-perfect match”) occurs if and only if:

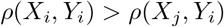

for all pairs (*i, j*) such that *i*≠ *j*.

Similarly, the event *M*_*X*_ (also called an “*X*-perfect match”) occurs if and only if

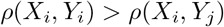

for all pairs (*i, j*) such that *i*≠ *j*.

#### Lemma 6

Let ***X*** = (*X*_1_, …, *X*_*n*_) be a sequence of random variables and let ***Y*** = (*Y*_1_, …, *Y*_*n*_) be a sequence of iid random variables. Let ***X*** and ***Y*** be independent. Then, the probability of a *Y*-perfect match is at most 1/*n*^*n*^.

**Proof:** Let ***X*** and ***Y*** be the sample spaces of ***X*** and ***Y*** (i.e. ***X*** ∈ ***X*** and ***Y*** ∈ ***Y***). Let ***c*** be a function that maps from ***X*** × ***Y*** to ℕ^*n*^. Specifically, ***c***(***x, y***) = (*c*_1_(***x, y***), …, *c*_*n*_(***x, y***)), where each *c*_*i*_ is given as follows:

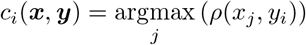

That is, *c*_*i*_(***x, y***) ∈ {1, …, *n*} is defined to be the index *j* that maximizes *ρ*(*x*_*j*_, *y*_*i*_). In the case of a multi-way tie, *c*_*i*_ is the lowest from among the maximizing index values. (For example, if *j* = 3 and *j* = 7 both produced the greatest value of *ρ*(*x*_*j*_, *y*_*i*_), then *c*_*i*_ would be 3 since 3 < 7.) Note that if a *Y*-perfect match occurs, then

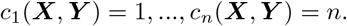

Consider ***c***(***x, Y***) for a fixed ***x*** ∈ ***X***. Notice that

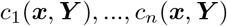

are iid because each *c*_*i*_(***x, Y***) depends only on *Y*_*i*_. Since the *c*_*i*_(***x, Y***) terms are identically distributed across *i*, we have:

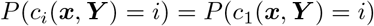

for all *i*. Therefore we have

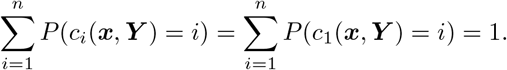

By dividing through by *n*, we may write

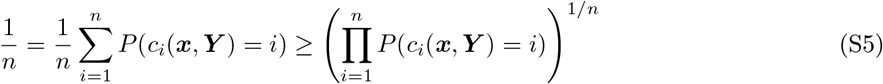

where the inequality comes from the AM-GM inequality (e.g., [43] pg. 20). Since the *c*_*i*_(***x, Y***) terms are independent,

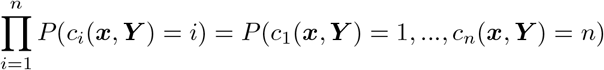

Substituting this fact into inequality S5 and raising both sides to the *n*th power gives us

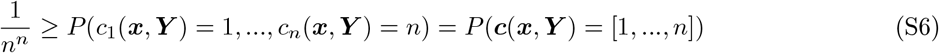

Since the above inequality holds for all ***x*** ∈ ***X***, and since ***X*** and ***Y*** are independent, Lemma 7 (given immediately following this proof) implies

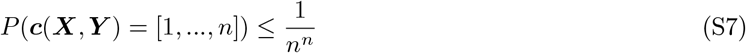

Since a *Y*-perfect match implies ***c***(***X, Y***) = [1, …, *n*], the left side of this inequality has a lower bound of *P* (*M*_*Y*_), giving us

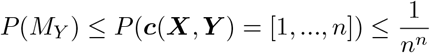

as required.

We now justify the step from Equation S6 to Equation S7 in the proof above. That is, we explain why a bound that holds for every fixed *x* continues to hold when *x* is replaced by the random variable *X*, provided *X* and *Y* are independent.

#### Lemma 7

Let *X* ∈ *X* and *Y* ∈ *Y* be independent random variables. Let *f* be a function whose domain is the product space *X* × *Y* and let *v* be a member of the range of *f*. Assume that there exists a constant *p* ∈ [0, 1] such that

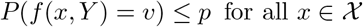

Then,

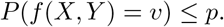

**Proof:** Let *I*_*v*_(*x, y*) denote the indicator function for the event where *f* (*x, y*) = *v*. That is, *I*_*v*_(*x, y*) = 1 if *f* (*x, y*) = *v* and otherwise *I*_*v*_(*x, y*) = 0.

From the law of total expectation we have

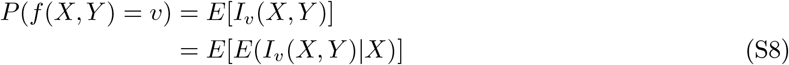

Let *g*(*x*) be a function given by

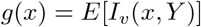

Since *X* and *Y* are independent, it follows (e.g., see Example 4.1.7 in [44]) that *g*(*X*) = *E*(*I*_*v*_(*X, Y*) *X*). Substituting this fact into equation S8 gives

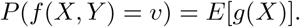

Note that it follows from the conditions of this lemma that *g*(*x*) ≤ *p* for all *x* ∈ X. Thus, monotonicity of the expected value implies *E*[*g*(*X*)] ≤ *E*[*p*] = *p*. Combining this fact with the above equation, we obtain

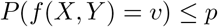

as required.

#### Lemma 8

Let ***X*** = (*X*_1_, …, *X*_*n*_) be a sequence of iid random variables and let ***Y*** = (*Y*_1_, …, *Y*_*n*_) be a sequence of random variables. Let ***X*** and ***Y*** be independent. Then, the probability of an *X*-perfect match is at most 1/*n*^*n*^.

**Proof:** The argument is analogous to that of Lemma 6. The main differences are that in this case we define 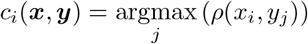 instead of *c*_*i*_(***x, y***) = argmax (*ρ*(*x*_*j*_, *y*_*i*_)), and here we condition on ***Y*** = ***y*** instead of ***X*** = ***x***.

#### Lemma 9

*A perfect match implies p*_*perm*_ = 1/*n*!

Let ***X*** = (*X*_1_, …, *X*_*n*_) and ***Y*** = (*Y*_1_, …, *Y*_*n*_) be sequences of random variables. Let *ρ* be a function that maps a pair (*X*_*i*_, *Y*_*j*_) to a real number. Define the average correlation

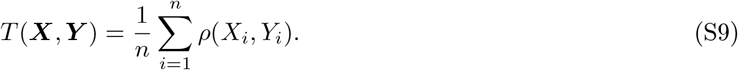

Perform the permutation test of independence (Def. 2) using *T* as the statistic, and obtain the *p*-value *p*_*perm*_. Also check for an *X*-perfect and *Y*-perfect match test (Def. 5) using *ρ* as the correlation function. Then, the occurrence of either an *X*-perfect match or a *Y*-perfect match implies that *p*_*perm*_ = 1/*n*!.

**Proof:** This claim is shown by a visual argument in Fig 1C. Here we walk through the same logic in prose.

Suppose a *Y*-perfect match occurs, meaning that

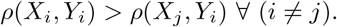

In other words, for each *Y*_*i*_, its correlation term *ρ*(, *Y*_*i*_) is strictly maximized when *Y*_*i*_ is paired with *X*_*i*_. Applying a permutation to ***Y*** (other than the trivial identity permutation) means that for at least one choice of *i*, the *ρ*(*X*_*i*_, *Y*_*i*_) term in Eq. S9 becomes replaced with *ρ*(*X*_*j*_, *Y*_*i*_) for some *j ≠ i*. But *ρ*(*X*_*i*_, *Y*_*i*_) >*ρ*(*X*_*j*_, *Y*_*i*_). Therefore, *T* (***X, Y***) must be strictly greater than all permuted versions *T* (***X***, *g*_*i*_***Y***) (other than when *g*_*i*_ is the identity permutation). It follows that *p*_*perm*_ = 1/*n*!.

Suppose now that an *X*-perfect match occurs. It follows that for each *X*_*i*_ its correlation term *ρ*(*X*_*i*_, ·) is strictly maximized when *X*_*i*_ is paired with *Y*_*i*_. Applying a permutation to ***X*** (other than the identity permutation) means that for at least one choice of *i*, the *ρ*(*X*_*i*_, *Y*_*i*_) term in Eq. S9 becomes replaced with *ρ*(*X*_*i*_, *Y*_*j*_) for some *j ≠ i*. Therefore, *T* (***X, Y***) is strictly greater than all permuted versions *T* (*g*_*i*_***X, Y***) whenever *g*_*i*_ is not the identity permutation. So *p*_*perm*_ = 1/*n*!.

#### Lemma 10

1/*n*! ≥ 2/*n*^*n*^ for all integers *n >* 1, with equality only when *n* = 2.

**Proof:** When *n* = 2 it is clear that 1/*n*! = 2/*n*^*n*^ = 1/2.

For *n >* 2, we establish 1/*n*! > 2/*n*^*n*^ by an inductive argument. For the base case, suppose *n* = 3. Then, 1/*n*! > 2/*n*^*n*^ becomes 1/6 > 2/27, which is true.

For the inductive step, suppose the target inequality 1/*n*! > 2/*n*^*n*^ holds. Multiplying the target inequality on both sides by 1/(*n* + 1), we obtain

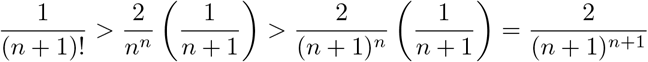

This shows that 1/*n*! > 2/*n*^*n*^ implies 1/(*n* + 1)! > 2/(*n* + 1)^*n*+1^. We conclude by the inductive principle that the target inequality is true for *n >* 2.

#### Theorem 11

*The permute-match*_*X*▶*Y*_ *test is valid*.

Let ***X*** = (*X*_1_, …, *X*_*n*_) and ***Y*** = (*Y*_1_, …, *Y*_*n*_) be sequences of iid random variables. Let ***X*** and ***Y*** be independent. Let the correlation function *ρ* be a function that maps a pair (*X*_*i*_, *Y*_*j*_) to a real number. Define the average correlation

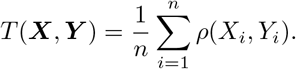

Perform the permutation test of independence (Def. 2) using *T* as the statistic, and obtain the *p*-value *p*_*perm*_. Also check for an *X*-perfect match test and a *Y*-perfect match test (Def. 5) using *ρ* as the correlation function. Let

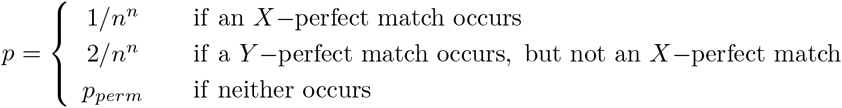

Then *P* (*p* ≤ *α*) ≤ *α* for *α* ∈ [0, 1] and *n >* 1.

**Proof:** For notational convenience, we denote the events of an *X*-perfect match and *Y*-perfect match as *M*_*X*_ and *M*_*Y*_.

There are four cases to consider. Either (case i) *α* ≥ 1/*n*!, or (case ii) 1/*n*! >*α* ≥ 2/*n*^*n*^, or (case iii) 2/*n*^*n*^ >*α* ≥ 1/*n*^*n*^, or (case iv) *α <* 1/*n*^*n*^. Note that Lemma 10 tells us that 1/*n*! ≥ 2/*n*^*n*^.

Case i: Suppose *α* ≥ 1/*n*!. In this case, *p* ≤ *α* if *M*_*X*_ occurs or *M*_*Y*_ occurs or *p*_*perm*_ ≤ *α*.

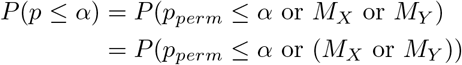

This may be split into 3 terms via the identity *P* (*A* or *B*) = *P* (*A* and *B*)+*P* (*A* and not *B*)+*P* ((not *A*) and *B*), as follows:

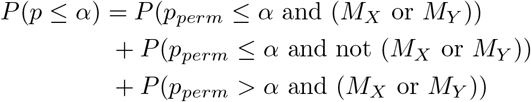

However, by Lemma 9, if either *M*_*X*_ or *M*_*Y*_ occurs, then *p*_*perm*_ = 1/*n*!, which would in turn imply *p*_*perm*_ ≤ *α* under case i. That is, the term *P* (*p*_*perm*_ >*α* and (*M*_*X*_ or *M*_*Y*_)) in the above summation is zero. Removing this term, we have

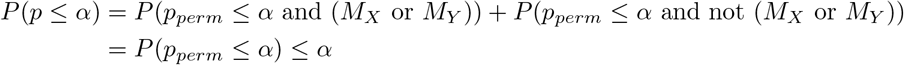

where the final inequality is due to Lemma 3. This establishes the claim for case i.

Case ii: Suppose 1/*n*! >*α* ≥ 2/*n*^*n*^. In this case, *p*_*perm*_ >*α* since *p*_*perm*_ cannot be less than 1/*n*!, so we have

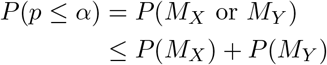

From Lemmas 6-8 we know *P* (*M*_*X*_) ≤ 1/*n*^*n*^ and *P* (*M*_*Y*_) ≤ 1/*n*^*n*^. Thus the above inequality becomes

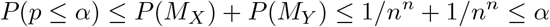

which establishes the claim in case ii.

Case iii: Suppose 2/*n*^*n*^ >*α* ≥ 1/*n*^*n*^. In this case, only *M*_*X*_ will give us *p* ≤ *α*. So we have

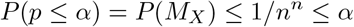

which verifies the claim in case iii.

Case iv: If *α <* 1/*n*^*n*^, then, *p* ≤ *α* is impossible so *P* (*p* ≤ *α*) = 0 ≤ *α*, so the claim is trivially satisfied in case iv.

Having been shown in all four cases, the claim has been proven.

#### Theorem 12

*The permute-match*_*Y* ▶*X*_ *test is valid*.

Consider the same setting as Theorem 11, except that now

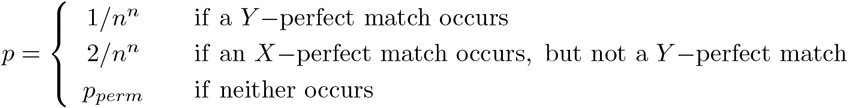

Then, *P* (*p* ≤ *α*) ≤ *α* for *α* ∈ [0, 1] and *n >* 1.

**Proof:** The proof is analogous to that of Theorem 11. In fact, all we need to change here is that in case iii, we use the fact that *P* (*M*_*Y*_) ≤ 1/*n*^*n*^ instead of that *P* (*M*_*X*_) ≤ 1/*n*^*n*^.

#### Theorem 13

*The simultaneous permute-match*_(*X*;*Y*)_ *test is valid*.

Consider the same setting as Theorem 11, except that now

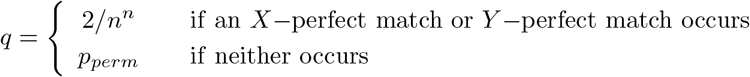

Then, *P* (*q* ≤ *α*) ≤ *α* for *α* ∈ [0, 1] and *n >* 1.

**Proof:** We first show by cases that *p* of Theorem 11 is always less than or equal to *q*. First, if an *X*-perfect match occurs, then *p* = 1/*n*^*n*^ < 2/*n*^*n*^ = *q*. Second, if there is a *Y*-perfect match but not an *X*-perfect match, then *p* = 2/*n*^*n*^ = *q*. Third, if neither an *X*-perfect nor a *Y*-perfect match occurs, then *p* = *p*_*perm*_ = *q*. Overall, *p* ≤ *q*. It follows that *q* ≤ *α* implies *p* ≤ *α*.

Then it follows from Theorem 11 that

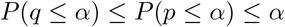

so *P* (*q* ≤ *α*) ≤ *α* as required.

We have seen that 1/*n*^*n*^ is a universal upper bound on *P* (*X* − perfect match). Here, we give an intuitive explanation for why this probability can be far smaller than 1/*n*^*n*^, and also why it is still possible to construct examples where *P* (*X* − perfect match) comes arbitrarily close to 1/*n*^*n*^.

To see why *P* (*X* − perfect match) can fall well below 1/*n*^*n*^, recall the analogy of students and advisors, where *ρ*(*X*_*i*_, *Y*_*j*_) measures how well a student *i* likes advisor *j* and an *X*-perfect match occurs if each student prefers their own advisor over all the others. Suppose that all the students value essentially the same qualities in an advisor. In the most extreme instance, *ρ*(*X*_*i*_, *Y*_*j*_) depends only on *Y*_*j*_ and not *X*_*i*_ at all. Then all students will rank advisors in the same way, so they will all prefer the same advisor, making an *X*-perfect match impossible.

This suggests that to maximize the chance of an *X*-perfect match, different students must care about different “dimensions” of their advisors. However, under the null hypothesis, the *X*_*i*_ variables are independent and identically distributed, so we cannot deterministically assign different advisor dimensions to different students. Instead, the key trick in our construction is to introduce an enormous number of possible dimensions. By allowing the number of advisor dimensions (*k* in the following formal construction) to be arbitrarily large, we make the probability that two students happen to care about the same dimension arbitrarily small. And, if indeed each student cares about a different (and independent) dimension, all the students will rank advisors independently, so that the probability of an *X*-perfect match approaches 1/*n*^*n*^.

The following formalizes the above intuition, establishing a scenario where under the null hypothesis, the probability of an *X*-perfect match can be made arbitrarily close to 1/*n*^*n*^. An analogous example could be constructed for a *Y*-perfect match.

**Construction** Let *k* be a positive integer whose value will be chosen later. For each *i* ∈ {1, …, *n*}, let *X*_*i*_ be drawn uniformly at random from among {1, …, *k*}, and let *X*_1_, …, *X*_*n*_ be independent.

For each *j* ∈ {1, …, *n*} and each *l* ∈ {1, …, *k*}, let

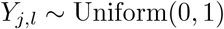

where all *Y*_*j,l*_ terms are independent of each other and of the *X*_*i*_ terms. Define *Y*_*j*_ = [*Y*_*j*,1_, …, *Y*_*j,k*_]. In the story above, *X*_*i*_ is the dimension that student *i* cares about, and *Y*_*j,l*_ is the quality of advisor *j* along dimension *l*.

Let the correlation function be

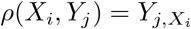

In this case, we say that an *X*-perfect match occurs if and only if

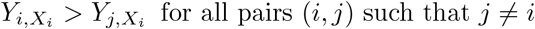

#### Proposition 14

Let *q* be a number such that 0 ≤ *q <* 1. Then, for the construction above, when *H*_0_ is satisfied (i.e., the *X*_*i*_ are iid, the *Y*_*i*_ are iid, and ***X*** and ***Y*** are independent), the probability of an *X*-perfect match is >*q/n*^*n*^.

**Proof:** Fix *x* ∈ {1, …, *k*} and let *A*_*j,x*_ be the event that *Y*_*j,x*_ is the strictly largest element from the sequence *Y*_[:,*x*]_ = [*Y*_1,*x*_, …, *Y*_*n,x*_]. Since the elements of *Y*_[:,*x*]_ are iid continuous random variables, the probability of a “tie” for largest is zero, and so it follows from symmetry that *P* (*A*_*j,x*_) = 1/*n* for all *j*. Now, fix {*x*_1_, …, *x*_*n*_} where each *x*_*i*_ is unique. Then, 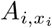 are all independent of one another because each 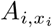 depends only on 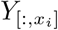, and the 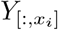 are independent. Thus

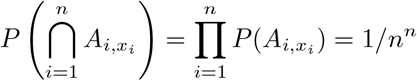

Notice that for ***X*** = ***x***, an *X*-perfect match occurs if and only if all the 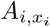 events occur. Let us formalize this further. Let M_*X*_ be a set in the product sample space of ***X*** and ***Y*** such that (***x, y***) ∈ M_*X*_ if and only if 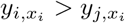 for all pairs *i*≠ *j*. Moreover, let us denote an *X*-perfect match by the symbol *M*_*X*_. Notice that *M*_*X*_ occurs if and only if (***X, Y***) ∈ M_*X*_. Now, we may rewrite the above equation as

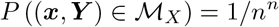

for all ***x*** whose entries are unique. Applying Lemma 15 (given just after the present proof) to the above result, we see that

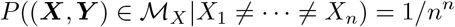

We are now ready to complete the result. We start with

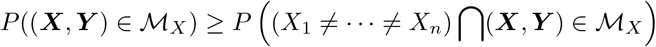

Substituting *M*_*X*_ ⇐⇒ (***X, Y***) ∈ M_*X*_, we now obtain:

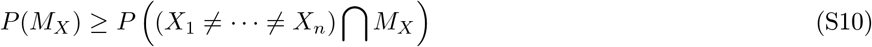

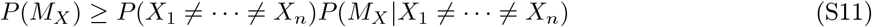

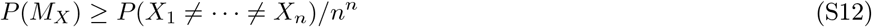

Since *X*_1_, …, *X*_*n*_ are uniformly distributed, we have

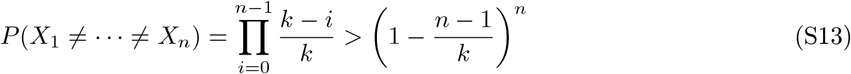

We have not yet specified *k*; we do so now. Choose *k* to be any integer with:

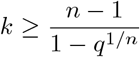

Then, combining this choice of *k* with inequality S13, we have *P* (*X*_1_ ≠ · · · ≠ *Xn*) > *q* substitute into inequality S12 to obtain *P* (*M*_*X*_) >*q/n*^*n*^, as required.

#### Lemma 15

Let *X* and *Y* be independent random variables. Let *A* be a set in the sample space of *X* with *P* (*X* ∈ *A*) > 0. Let *B* be a set in the product space of *X* and *Y*. Assume that there exists a constant *p* ∈ [0, 1] such that

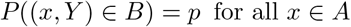

Then,

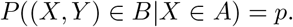

**Proof:** Let *I*_*B*_(*x, y*) denote the indicator function for the event where (*x, y*) ∈ *B*. That is, *I*_*B*_(*x, y*) = 1 if (*x, y*) ∈ *B* and otherwise *I*_*B*_(*x, y*) = 0. Let *g*(*x*) be a function given by

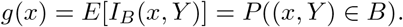

Note that it follows from the conditions of this lemma that *g*(*x*) = *p* for all *x* ∈ *A*. Consider the joint probability

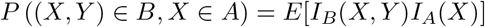

where *I*_*A*_(*x*) is the indicator function for *x* ∈ *A*. From the law of total expectation, the above is equal to

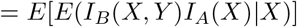

since *I*_*A*_(*X*) is a function of *X* only, we may pull it out of the inner conditional expectation (“taking out what is known”; see for instance Theorem 9.3.2 in [45]):

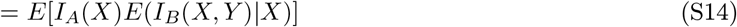

Since *X* and *Y* are independent, it follows from Example 4.1.7 in [44] that *g*(*X*) = *E*(*I*_*B*_(*X, Y*) | *X*).

Substituting this result into Eq. S14 gives

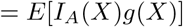

Notice that *I*_*A*_(*x*)*g*(*x*) = *I*_*A*_(*x*)*p* because both evaluate to *p* for all *x* ∈ *A*, and evaluate to 0 for all *x /*∈ *A*. Thus the above becomes

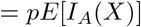

To summarize this chain of equalities, we now have

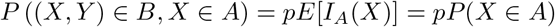

Finally, by substituting this result into the definition of conditional probability, we have

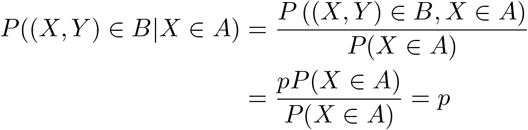

as required.

### 3 The sequential permute-match test versus a seemingly similar flawed procedure

In the main text we briefly noted that the sequential permute-match test superficially resembles the flawed practice of conducting additional tests merely because earlier tests did not result in a detection. Here we provide a detailed explanation of why this practice generally results in an invalid *p*-value, and point out that the sequential permute-match test does not have this problem.

To spell out the flawed practice, suppose a researcher runs one test (test *X*) and obtains a *p*-value, *p*_*X*_. If *p*_*X*_ is below their desired significance level *α*, they stop there and report *p*_*X*_. Conversely, if *p*_*X*_ >*α*, this researcher performs another test (test *Y*) that assesses the same biological hypothesis in a slightly different way, obtaining a second *p*-value, *p*_*Y*_. However, the researcher has a vague understanding that if one conducts multiple tests, a correction is required, so at this point they perform a Bonferroni correction to account for two tests and report 2*p*_*Y*_. Overall, the *p*-value obtained from this series of steps is:

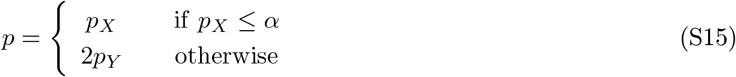

One may ask whether this overall procedure has the desired property (known as ‘validity’) that it prevents the false positive rate from exceeding the significance level. Unfortunately, this procedure is generally not valid, meaning that it commonly leads to *P* (*p* ≤ *α*) >*α* under the null hypothesis.

To demonstrate this fact mathematically, we make three assumptions. First, we assume that the null hypothesis of independence is true for both tests *X* and *Y* so that *P* (*p* ≤ *α*) is the false positive rate. Second, we assume that *p*_*X*_ and *p*_*Y*_ are both random variables following a uniform distribution between 0 and 1. Finally, we assume that the significance level *α* is less than 1/2.

The false positive rate of the procedure in Eq S15 depends on the relationship between the two tests. If tests *X* and *Y* tend to report the same result, they will behave more like a single test and the false positive rate of Eq S15 will be lower; if the tests frequently report different results, they will behave like two genuinely distinct tests, and the false positive rate will be greater. This relationship is quantified in Table S1, which describes the joint and marginal distributions of the *X* and *Y* test outcomes. We write the probability that test *X* is significant (or not) along the bottom row, and the probability that test *Y* is significant (or not) along the rightmost column. We then define a parameter (*P*′) which is the probability that test *Y* is significant while test *X* is not. We then fill out the rest of the table, corresponding to the probabilities of the other possible test results, by ensuring that the rows and columns of the 2 × 2 “Joint probability” table sum to the entries in the “Total probability” row and column.

**Table S1:** Joint and marginal probabilities of the outcomes of tests *X* and *Y*, assuming that *p*_*X*_ and *p*_*Y*_ both follow Unif(0, 1). We first fill the total probability row and column, and define *P*′ as the probability that test *Y* is significant but test *X* is not. The remaining entries can then be deduced.

|  |  | Joint probability |  | Total probability |
| --- | --- | --- | --- | --- |
| | | Test $X$ is significant<br>($p_X \leq \alpha$) | Test $X$ not significant<br>($p_X > \alpha$) | |
| Joint probability | Test $Y$ not significant<br>($2p_Y > \alpha$) | $\alpha/2 + P'$ | $1 - \alpha - P'$ | $1 - \alpha/2$ |
| | Test $Y$ is significant<br>($2p_Y \leq \alpha$) | $\alpha/2 - P'$ | $P'$ | $\alpha/2$ |
| Total probability | | $\alpha$ | $1 - \alpha$ | |

Each joint probability is of course bounded between 0 and 1. Thus, moving clockwise from the upper left of the 2 × 2 inner table we have the following constraints on *P*′:

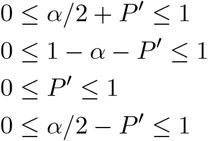

Among these constraints, since *α <* 1/2, the tightest bounds on *P*′ are 0 ≤ *P*′ ≤ *α/*2. Notice that the false positive rate of the overall procedure is given by:

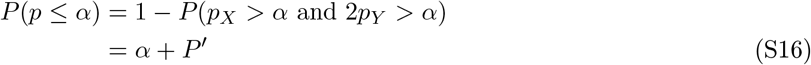

Fig S1 plots Eq S16 within the allowed bounds of *P*′, noting the extreme cases of *P*′ = 0 and *P*′ = *α/*2, as well as the case of independence in which *P*′ = *P* (2*p*_*Y*_ ≤ *α*) × (*p*_*X*_ >*α*). Other than the extreme case of *P*′ = 0, in which a significant *Y* test result deterministically ensures that the *X* test is also significant, the false positive rate of the overall procedure, *P* (*p* ≤ *α*), will be greater than *α*. Thus the procedure is not valid.

The approach for the sequential permute-match test (Fig 1F) is different because it is based on discrete events, not continuous random variables. Perhaps counterintuitively, we can actually get away with a similar recipe while producing a valid *p*-value. To best compare our approach to the flawed procedure above, we presently leave out the permutation part and focus only on the matching test component in the following proposition:

#### Proposition 16

Let *M*_*X*_ and *M*_*Y*_ be events with *P* (*M*_*X*_) ≤ 1/*n*^*n*^ and *P* (*M*_*Y*_) ≤ 1/*n*^*n*^. Let

**Figure S1:**
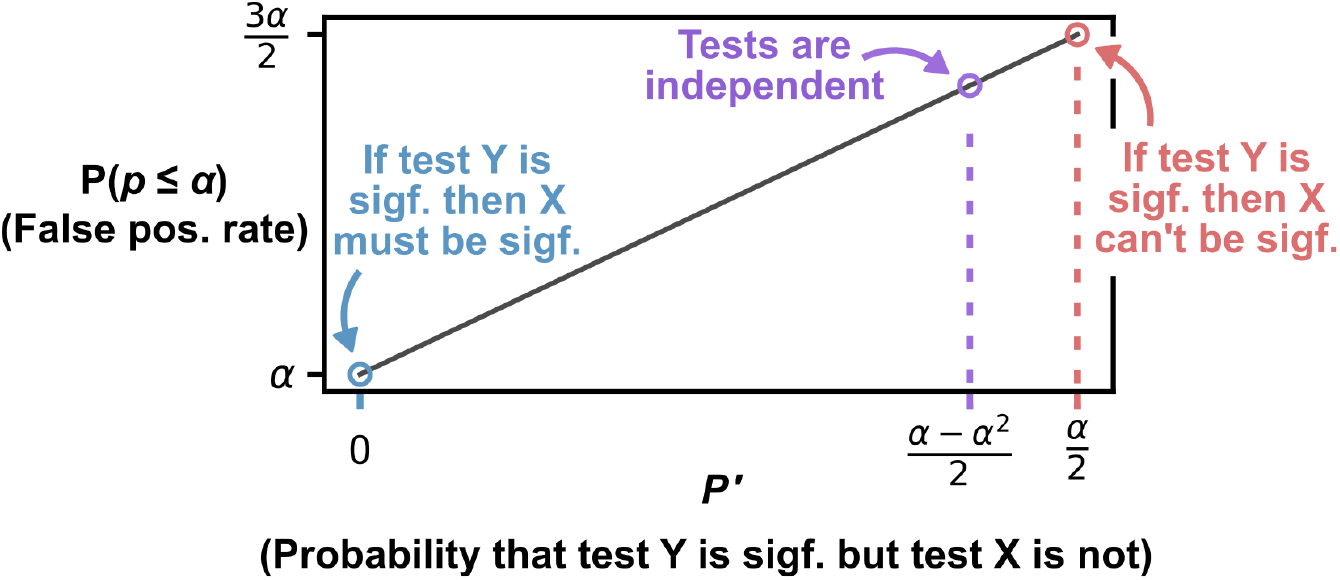
The procedure of Eq S15 produces a false positive rate that generally exceeds *α*, and depends on the relationship between tests *X* and *Y*. This relationship between *X* and *Y* can be quantified by *P*′, the probability that test *Y* is significant (i.e. 2*p*_*Y*_ ≤ *α*) but test *X* is not significant (i.e. *p*_*X*_ >*α*). The false positive rate is shown over the full range of *P*′, from 0 to *α/*2. The only way for the overall procedure of Eq S15 to be valid is when *P*′ = 0, where tests *X* and *Y* are so tightly coupled that a significant result in test *Y* deterministically ensures that test *X* is also significant. Other than this edge case, the false positive rate of the overall procedure will exceed *α*.

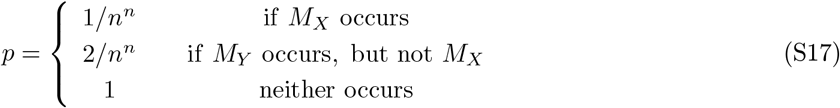

Then, *P* (*p* ≤ *α*) ≤ *α* for 0 ≤ *α* ≤ 1 and for all integers *n* ≥ 2.

**Proof:** We consider four cases, each corresponding to possible values of *α*.

First, if *α* = 1, then *P* (*p* ≤ *α*) ≤ *α* trivially because probability does not exceed 1.

Second is the case 1 >*α* ≥ 2/*n*^*n*^. In this case, either *M*_*X*_ or *M*_*Y*_ is sufficient to achieve *p* ≤ *α*. So we have:

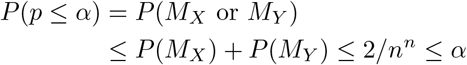

as required.

Third, consider the case 2/*n*^*n*^ >*α* ≥ 1/*n*^*n*^. In this case, only *M*_*X*_ will give us *p* ≤ *α*. So we have

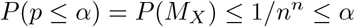

as required.

Finally consider the case *α <* 1/*n*^*n*^. In this case *p* ≤ *α* is impossible so *P* (*p* ≤ *α*) = 0 ≤ *α*, as required.

Since the claim holds under our (exhaustive) set of cases, the proof is complete.

The same argument holds if 1/*n*^*n*^ and 2/*n*^*n*^ are replaced by *q* and 2*q* for an arbitrary *q* between 0 and 1, but we use 1/*n*^*n*^ and 2/*n*^*n*^ to highlight the connection to the permute-match test. Our complete proof (Theorem 11) followed largely the same steps as shown here. Overall, we hope that readers of this section now have a deeper understanding of why the sequential permute-match test is valid, and an appreciation of why similar approaches are invalid in the more standard context where statistical tests are not simply binary event checks.

### 4 Illustration of permute-match test with logistic map system

**Figure S2:**
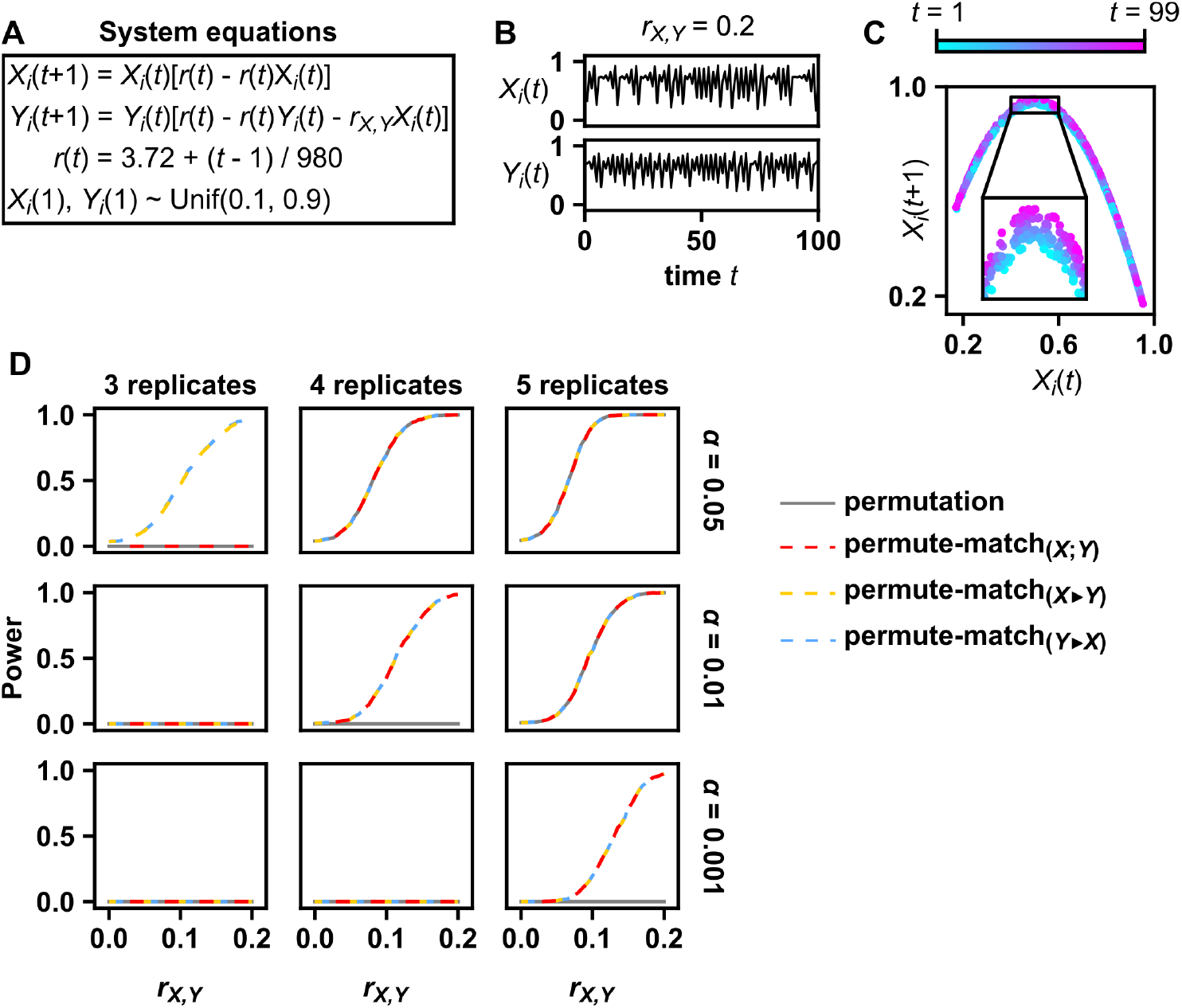
Statistical power of permute-match tests and the permutation test in a nonlinear and nonstationary system. (**A**) System equations. (**B**) Example dynamics. The processes *X* and *Y* are given by a coupled logistic map on a time grid *t* = 1, 2, …, 100. The system is nonstationary because the parameter *r*(*t*) varies with time from 3.72 when *t* = 1 to 3.82 when *t* = 99. (**C**) To see the nonstationarity clearly, we plot *X*_*i*_(*t*) against *X*_*i*_(*t* + 1) and color points by time, showing that the parabola that the points lie on is drifting over time. To better show the trend, 10 replicates are shown simultaneously in this chart. (**D**) Statistical power of the permutation test and permute-match tests as a function of the number of replicates, the significance level, and *r*_*X,Y*_. Power was estimated from 5000 simulations at each value of *r*_*X,Y*_ between *r*_*X,Y*_ = 0 and *r*_*X,Y*_ = 0.2 in steps of size 0.005. The correlation statistic (*ρ*) was cross-map skill, which is known to readily detect dependence between *X* and *Y* in this system [46]. For the cross-map skill calculation, we used *Y* to estimate *X*, which corresponds to a scenario where the data analyst hypothesizes that *X* influences *Y*. Cross-map skill requires two parameters, the embedding dimension and the embedding lag, and these were set to 2 and 1 respectively following prior works that used the logistic map for benchmarking [46, 47].

**Figure S3:**
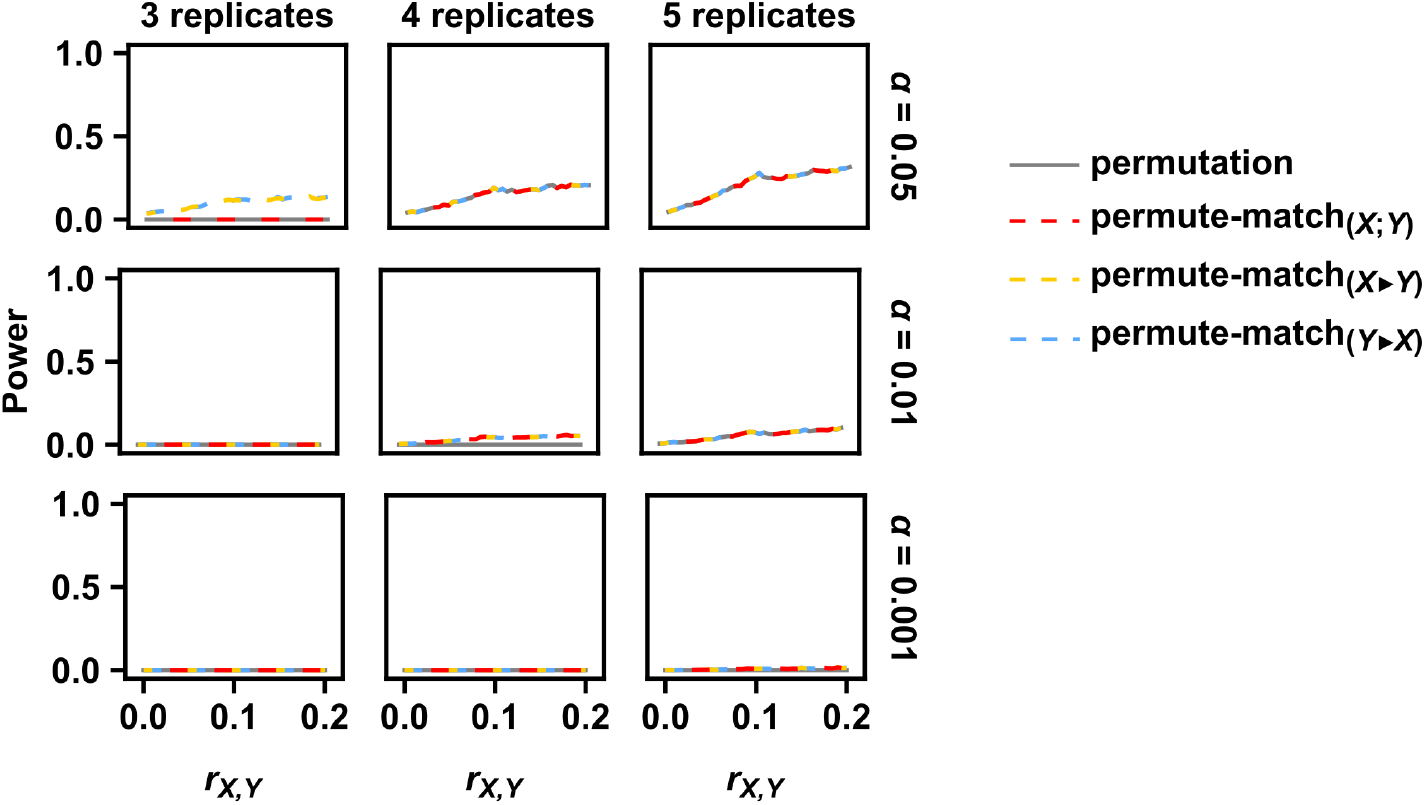
Pearson correlation has lower power than cross-map skill for the logistic map. Shown here is the same result as in Fig S2D, except now the correlation statistic (*ρ*) is Pearson correlation as in Fig 2.

