## Supplementary material for "Permute-match tests detect significant correlations between time series despite nonstationarity and limited replicates": Source Code 1: local.yml

```
name: uw_mac
channels:
  - plotly
  - conda-forge
  - defaults
dependencies:
  - _ipyw_jlab_nb_ext_conf=0.1.0=py37_0
  - alabaster=0.7.12=py37_0
  - anaconda=2019.10=py37_0
  - anaconda-client=1.7.2=py37_0
  - anaconda-navigator=1.9.7=py37_0
  - anaconda-project=0.8.3=py_0
  - appnope=0.1.0=py37_0
  - appscript=1.1.0=py37h1de35cc_0
  - asn1crypto=1.0.1=py37_0
  - astroid=2.3.1=py37_0
  - astropy=3.2.2=py37h1de35cc_0
  - atomicwrites=1.3.0=py37_1
  - attrs=19.2.0=py_0
  - babel=2.7.0=py_0
  - backcall=0.1.0=py37_0
  - backports=1.0=py_2
  - backports.functools_lru_cache=1.6.4=pyhd3eb1b0_0
  - backports.os=0.1.1=py37_0
  - backports.shutil_get_terminal_size=1.0.0=py37_2
  - backports.tempfile=1.0=pyhd3eb1b0_1
  - backports.weakref=1.0.post1=py_1
  - beautifulsoup4=4.8.0=py37_0
  - bitarray=1.0.1=py37h1de35cc_0
  - bkcharts=0.2=py37_0
  - blas=1.0=mkl
  - bleach=3.1.0=py37_0
  - blosc=1.16.3=hd9629dc_0
  - bokeh=1.3.4=py37_0
  - boto=2.49.0=py37_0
  - bottleneck=1.2.1=py37h1d22016_1
  - bzip2=1.0.8=h1de35cc_0
  - ca-certificates=2019.8.28=0
  - certifi=2019.9.11=py37_0
  - cffi=1.12.3=py37hb5b8e2f_0
  - chardet=3.0.4=py37_1003
  - click=7.0=py37_0
  - cloudpickle=1.2.2=py_0
  - clyent=1.2.2=py37_1
  - colorama=0.4.1=py37_0
  - conda-package-handling=1.7.3=py37h9ed2024_1
  - conda-verify=3.4.2=py_1
  - contextlib2=0.6.0=py_0
  - cryptography=2.7=py37ha12b0ac_0
  - curl=7.65.3=ha441bb4_0
  - cycler=0.10.0=py37_0
  - cython=0.29.13=py37h0a44026_0
  - cytoolz=0.10.0=py37h1de35cc_0
  - dask=2.5.2=py_0
  - dask-core=2.5.2=py_0
  - dbus=1.13.6=h90a0687_0
  - decorator=4.4.0=py37_1
  - defusedxml=0.6.0=py_0
  - distributed=2.5.2=py_0
  - docutils=0.15.2=py37_0
  - entrypoints=0.3=py37_0
  - et_xmlfile=1.0.1=py37_0
  - expat=2.2.6=h0a44026_0
  - fastcache=1.1.0=py37h1de35cc_0
  - ffmpeg=4.0=h01ea3c9_0
  - flask=1.1.1=py_0
  - freetype=2.9.1=hb4e5f40_0
  - fsspec=0.5.2=py_0
  - future=0.18.2=py37_1
  - get_terminal_size=1.0.0=h7520d66_0
  - gettext=0.19.8.1=h15daf44_3
  - gevent=1.4.0=py37h1de35cc_0
  - glib=2.56.2=hd9629dc_0
  - glob2=0.7=py_0
  - gmp=6.1.2=hb37e062_1
  - gmpy2=2.0.8=py37h6ef4df4_2
  - greenlet=0.4.15=py37h1de35cc_0
  - h5py=2.9.0=py37h3134771_0
  - hdf5=1.10.4=hfa1e0ec_0
  - heapdict=1.0.1=py_0
  - html5lib=1.0.1=py37_0
  - icu=58.2=h4b95b61_1
  - idna=2.8=py37_0
  - imageio=2.6.0=py37_0
  - imagesize=1.1.0=py37_0
  - intel-openmp=2019.4=233
  - ipykernel=5.1.2=py37h39e3cac_0
  - ipython=7.8.0=py37h39e3cac_0
  - ipython_genutils=0.2.0=py37_0
  - ipywidgets=7.5.1=py_0
  - isort=4.3.21=py37_0
  - itsdangerous=1.1.0=py37_0
  - jbig=2.1=h4d881f8_0
  - jdcal=1.4.1=py_0
  - jedi=0.15.1=py37_0
  - jinja2=2.10.3=py_0
  - jpeg=9b=he5867d9_2
  - json5=0.8.5=py_0
  - jsonschema=3.0.2=py37_0
  - jupyter=1.0.0=py37_7
  - jupyter_client=5.3.3=py37_1
  - jupyter_console=6.0.0=py37_0
  - jupyter_core=4.5.0=py_0
  - jupyterlab=1.1.4=pyhf63ae98_0
  - jupyterlab_server=1.0.6=py_0
  - keyring=18.0.0=py37_0
  - kiwisolver=1.1.0=py37h0a44026_0
  - krb5=1.16.1=hddcf347_7
  - lazy-object-proxy=1.4.2=py37h1de35cc_0
  - libarchive=3.3.3=h786848e_5
  - libcurl=7.65.3=h051b688_0
  - libcxx=4.0.1=hcfea43d_1
  - libcxxabi=4.0.1=hcfea43d_1
  - libedit=3.1.20181209=hb402a30_0
  - libffi=3.2.1=h475c297_4
  - libgfortran=3.0.1=h93005f0_2
  - libiconv=1.15=hdd342a3_7
  - liblief=0.9.0=h2a1bed3_2
  - libopus=1.3.1=h1de35cc_0
  - libpng=1.6.37=ha441bb4_0
  - libsodium=1.0.16=h3efe00b_0
  - libssh2=1.8.2=ha12b0ac_0
  - libtiff=4.0.10=hcb84e12_2
  - libvpx=1.7.0=h378b8a2_0
  - libxml2=2.9.9=hf6e021a_1
  - libxslt=1.1.33=h33a18ac_0
  - llvm-openmp=4.0.1=hcfea43d_1
  - llvmlite=0.29.0=py37h98b8051_0
  - locket=0.2.0=py37_1
  - lxml=4.4.1=py37hef8c89e_0
  - lz4-c=1.8.1.2=h1de35cc_0
  - lzo=2.10=h362108e_2
  - markupsafe=1.1.1=py37h1de35cc_0
  - mccabe=0.6.1=py37_1
  - mistune=0.8.4=py37h1de35cc_0
  - mkl=2019.4=233
  - mkl-service=2.3.0=py37hfbe908c_0
  - mkl_fft=1.0.14=py37h5e564d8_0
  - mkl_random=1.1.0=py37ha771720_0
  - mock=3.0.5=py37_0
  - more-itertools=7.2.0=py37_0
  - mpc=1.1.0=h6ef4df4_1
  - mpfr=4.0.1=h3018a27_3
  - mpmath=1.1.0=py37_0
  - msgpack-python=0.6.1=py37h04f5b5a_1
  - multipledispatch=0.6.0=py37_0
  - navigator-updater=0.2.1=py37_0
  - nbconvert=5.6.0=py37_1
  - nbformat=4.4.0=py37_0
  - ncurses=6.1=h0a44026_1
  - networkx=2.3=py_0
  - nltk=3.4.5=py37_0
  - nose=1.3.7=py37_2
  - notebook=6.0.1=py37_0
  - numba=0.45.1=py37h6440ff4_0
  - numexpr=2.7.0=py37h7413580_0
  - numpydoc=0.9.1=py_0
  - olefile=0.46=py37_0
  - openpyxl=3.0.0=py_0
  - openssl=1.1.1d=h1de35cc_2
  - packaging=19.2=py_0
  - pandas=0.25.1=py37h0a44026_0
  - pandoc=2.2.3.2=0
  - pandocfilters=1.4.2=py37_1
  - parso=0.5.1=py_0
  - partd=1.0.0=py_0
  - path.py=12.0.1=py_0
  - pathlib2=2.3.5=py37_0
  - patsy=0.5.1=py37_0
  - pcre=8.43=h0a44026_0
  - pep8=1.7.1=py37_0
  - pexpect=4.7.0=py37_0
  - pickleshare=0.7.5=py37_0
  - pillow=6.2.0=py37hb68e598_0
  - pip=19.2.3=py37_0
  - pkginfo=1.5.0.1=py37_0
  - plotly=4.9.0=py_0
  - pluggy=0.13.0=py37_0
  - ply=3.11=py37_0
  - prometheus_client=0.7.1=py_0
  - prompt_toolkit=2.0.10=py_0
  - psutil=5.6.3=py37h1de35cc_0
  - ptyprocess=0.6.0=py37_0
  - py=1.8.0=py37_0
  - py-lief=0.9.0=py37h1413db1_2
  - pycodestyle=2.5.0=py37_0
  - pycosat=0.6.3=py37h1de35cc_0
  - pycparser=2.19=py37_0
  - pycrypto=2.6.1=py37h1de35cc_9
  - pycurl=7.43.0.3=py37ha12b0ac_0
  - pyflakes=2.1.1=py37_0
  - pygments=2.4.2=py_0
  - pylint=2.4.2=py37_0
  - pyodbc=4.0.27=py37h0a44026_0
  - pyopenssl=19.0.0=py37_0
  - pyparsing=2.4.2=py_0
  - pyqt=5.9.2=py37h655552a_2
  - pyrsistent=0.15.4=py37h1de35cc_0
  - pysocks=1.7.1=py37_0
  - pytables=3.5.2=py37h5bccee9_1
  - pytest=5.2.1=py37_0
  - pytest-arraydiff=0.3=py37h39e3cac_0
  - pytest-astropy=0.5.0=py37_0
  - pytest-doctestplus=0.4.0=py_0
  - pytest-openfiles=0.4.0=py_0
  - pytest-remotedata=0.3.2=py37_0
  - python=3.7.4=h359304d_1
  - python-dateutil=2.8.0=py37_0
  - python-libarchive-c=2.8=py37_13
  - python.app=2=py37_9
  - pytz=2019.3=py_0
  - pywavelets=1.0.3=py37h1d22016_1
  - pyyaml=5.1.2=py37h1de35cc_0
  - pyzmq=18.1.0=py37h0a44026_0
  - qt=5.9.7=h468cd18_1
  - qtawesome=0.6.0=py_0
  - qtconsole=4.5.5=py_0
  - qtpy=1.9.0=py_0
  - readline=7.0=h1de35cc_5
  - requests=2.22.0=py37_0
  - retrying=1.3.3=py37_2
  - ripgrep=0.10.0=hc07d326_0
  - rope=0.14.0=py_0
  - ruamel_yaml=0.15.46=py37h1de35cc_0
  - scikit-image=0.15.0=py37h0a44026_0
  - seaborn=0.9.0=py37_0
  - send2trash=1.5.0=py37_0
  - setuptools=41.4.0=py37_0
  - simplegeneric=0.8.1=py37_2
  - singledispatch=3.4.0.3=py37_0
  - sip=4.19.8=py37h0a44026_0
  - six=1.12.0=py37_0
  - snappy=1.1.7=he62c110_3
  - snowballstemmer=2.0.0=py_0
  - sortedcollections=1.1.2=py37_0
  - sortedcontainers=2.1.0=py37_0
  - soupsieve=1.9.3=py37_0
  - sphinx=2.2.0=py_0
  - sphinxcontrib=1.0=py37_1
  - sphinxcontrib-applehelp=1.0.1=py_0
  - sphinxcontrib-devhelp=1.0.1=py_0
  - sphinxcontrib-htmlhelp=1.0.2=py_0
  - sphinxcontrib-jsmath=1.0.1=py_0
  - sphinxcontrib-qthelp=1.0.2=py_0
  - sphinxcontrib-serializinghtml=1.1.3=py_0
  - sphinxcontrib-websupport=1.1.2=py_0
  - spyder=3.3.6=py37_0
  - spyder-kernels=0.5.2=py37_0
  - sqlalchemy=1.3.9=py37h1de35cc_0
  - sqlite=3.30.0=ha441bb4_0
  - statsmodels=0.10.1=py37h1d22016_0
  - sympy=1.4=py37_0
  - tbb=2019.8=h04f5b5a_0
  - tblib=1.4.0=py_0
  - terminado=0.8.2=py37_0
  - testpath=0.4.2=py37_0
  - tk=8.6.8=ha441bb4_0
  - toolz=0.10.0=py_0
  - tornado=6.0.3=py37h1de35cc_0
  - tqdm=4.36.1=py_0
  - traitlets=4.3.3=py37_0
  - unicodecsv=0.14.1=py37_0
  - unixodbc=2.3.7=h1de35cc_0
  - urllib3=1.24.2=py37_0
  - wcwidth=0.1.7=py37_0
  - webencodings=0.5.1=py37_1
  - werkzeug=0.16.0=py_0
  - wheel=0.33.6=py37_0
  - widgetsnbextension=3.5.1=py37_0
  - wrapt=1.11.2=py37h1de35cc_0
  - wurlitzer=1.0.3=py37_0
  - xlrd=1.2.0=py37_0
  - xlsxwriter=1.2.1=py_0
  - xlwings=0.15.10=py37_0
  - xlwt=1.3.0=py37_0
  - xz=5.2.4=h1de35cc_4
  - yaml=0.1.7=hc338f04_2
  - zeromq=4.3.1=h0a44026_3
  - zict=1.0.0=py_0
  - zipp=0.6.0=py_0
  - zlib=1.2.11=h1de35cc_3
  - zstd=1.3.7=h5bba6e5_0
  - pip:
      - biom-format==2.1.10
      - conda-pack==0.6.0
      - cvxpy==1.2.0
      - distlib==0.3.6
      - ecos==2.0.10
      - filelock==3.8.0
      - giotto-tda==0.5.1
      - importlib-metadata==4.12.0
      - install==1.3.5
      - joblib==1.1.0
      - kdepy==1.1.0
      - matlabruntimeforpython==R2022a
      - matplotlib==3.4.0
      - miniballcpp==0.2.2
      - numpy==1.21.6
      - osqp==0.6.2.post5
      - platformdirs==2.5.2
      - pybind11==2.5.0
      - pyedm==1.3.7.1
      - pyflagser==0.4.4
      - python-igraph==0.8.2
      - pyunicorn==0.6.1
      - qdldl==0.1.5.post0
      - scikit-learn==1.0.2
      - scipy==1.7.3
      - scs==3.2.0
      - selenium==3.141.0
      - texttable==1.6.2
      - threadpoolctl==3.1.0
      - trajectorytools==0.3.4a0
      - typing-extensions==4.3.0
      - virtualenv==20.16.3
```
