## Supplementary material for "Permute-match tests detect significant correlations between time series despite nonstationarity and limited replicates": Source Code 1: fish_example.py

```
# Analyze fish data using permute-match tests

# Notes:
# ignoring nans for speed and var calc; ignoring still fish for var calc.
# speed in units of BL / sec

import numpy as np
import pandas as pd
import seaborn as sns
import os
import sys
import matplotlib.pyplot as plt
sys.path.append('../main_benchmark')
from examples import match_test, corrcoef, improved_permtest
from matplotlib import rc, rcParams
from scipy import stats
from statsmodels.tsa.stattools import kpss
from scipy.stats import pearsonr

# matplotlib settings
plt.rcParams['svg.fonttype'] = 'none'
rcParams['axes.linewidth'] = 1.5
rcParams['xtick.major.width'] = 1.5
rcParams['xtick.major.size'] = 5
rcParams['ytick.major.width'] = 1.5
rcParams['ytick.major.size'] = 5

font = {'family' : 'Arial',
        'weight' : 'bold',
        'size'   : 12}

rc('font', **font)

def get_position_and_velocity(trajectories_path, verbose=True):
    # reports position in units of fish lengths
    # and velocity in units of units of fish lengths per second
    data = np.load(trajectories_path, allow_pickle=True, encoding='bytes')
    position = data.tolist()['trajectories']
    framerate = data.tolist()['frames_per_second']
    body_length_px = 1 / data.tolist()['body_length']
    if verbose:
        print(f'frames per second: {framerate}')
        print(f'body length in pixels: {1/body_length_px}')
    # compute velocity via simple 2-point method
    velocity = (position[2:,:,:] - position[:-2,:,:]) / 2
    # truncate position to make dims consistent
    position = position[1:-1,:,:]
    # unit conversion
    position *= body_length_px
    velocity *= body_length_px * framerate
    return position, velocity

def load_all_data():
    path_to_data_folder = '/Users/alexyuan/Documents/Shou Lab/timeshift_properties/timeshift_repo/real_data/data/idtracker_ai/zebrafish_trajectories/10/' # using small groups of zebrafish
    position_all = []
    velocity_all = []
    min_ts_len = np.inf
    for experiment_id in [1,2,3]:
        trajectories_path = os.path.join(path_to_data_folder, str(experiment_id), 'trajectories_wo_gaps.npy')
        position, velocity = get_position_and_velocity(trajectories_path, verbose=False)
        position_all.append(position)
        velocity_all.append(velocity)
        min_ts_len = np.min([min_ts_len, position.shape[0]]).astype(int)
        print(f'length of velocity time series in trial {experiment_id}: {velocity.shape[0]} frames = {velocity.shape[0] / 32 / 60} minutes') # number of velocity frames is total number of frames - 2
    position_all = np.concatenate([
    np.expand_dims(_[:min_ts_len,:,:], axis=-1) for _ in position_all
    ], axis=-1)
    velocity_all = np.concatenate([
    np.expand_dims(_[:min_ts_len,:,:], axis=-1) for _ in velocity_all
    ], axis=-1)
    return position_all, velocity_all


def kpss_helper(
    timeseries, # 1d time series
    variable, # variable label, such as 'avg spd mvg avg'
    trial, # trial: 1,2, or 3
    kpss_results, # dict we're appending
    nan_policy = 'strict', # or 'exclude'
    **kpss_kwargs,
    ):
    if nan_policy == 'strict':
        assert np.all(np.isfinite(timeseries)), "nan_policy is strict: all values in timeseries must be finite."
        kpss_output = kpss(timeseries, **kpss_kwargs)
    elif nan_policy == 'exclude':
        finite_idx = np.isfinite(timeseries)
        kpss_output = kpss(timeseries[finite_idx], **kpss_kwargs)
    else:
        raise ValueError('nan_policy must be strict or exclude.')

    this_kpss_result = {
        'variable' : variable,
        'trial' : trial,
        'p<=' : kpss_output[1],
        '1% crit val' : kpss_output[3]['1%'],
        'statistic' : kpss_output[0]
    }
    for (k, v) in this_kpss_result.items():
        kpss_results[k].append(v)

position, velocity = load_all_data()
# now position and velocity have dims:
# frame (0,...,19308)
# fish (0,...,9)
# dimension (x, y)
# trial (1,2,3)

# investigate nans
speed = (velocity[:,:,0,:]**2 + velocity[:,:,1,:]**2)**(1/2)

print('number of still fish-frames per trial')
print(np.sum(np.sum(speed==0, axis=0), axis=0))

good_data = np.ones(speed.shape)
good_data[speed == 0] = 0 # cannot define direction of motion when speed = 0
good_data[np.logical_not(np.all(np.isfinite(velocity), axis=2))] = 0 # if either x or y data are bad, the whole fish-frame is unusable

print('number and percentage of bad fish-frames per trial')
print(np.sum(np.sum(1-good_data, axis=0),axis=0))
print(np.mean(np.mean(1-good_data, axis=0),axis=0))

print('fewest usable fish across all frames and trials')
print(np.min(good_data.sum(axis=1)))

avg_speed = np.nanmean(speed, axis=1)

velocity_uvecs = velocity / np.expand_dims(speed, axis=2)
good_data_broadcast = np.concatenate(
    [np.expand_dims(good_data,2), np.expand_dims(good_data,2)], axis=2
) # still a 1 if both x and y are good and 0 otherwise
for xy in [0,1]:
    assert np.isfinite(velocity_uvecs[good_data_broadcast.astype('bool')]).all()
    velocity_uvecs[np.logical_not(good_data_broadcast.astype('bool'))] = 0 # bad data excluded from sum

sum_velocity_uvecs = np.sum(velocity_uvecs, axis=1) # sum of unit vectors from good data
total_resultant_length = (sum_velocity_uvecs[:,0,:]**2 + sum_velocity_uvecs[:,1,:]**2)**(1/2)
mean_resultant_length = total_resultant_length / good_data.sum(axis=1) # divide by number of good-data fish

circ_conc = mean_resultant_length # rename to align with old code
print(f'number of frames in shortest time series: {circ_conc.shape[0]}')
print(f'in minutes: {circ_conc.shape[0] / 32 / 60}')

# perform the perfect match test; store results
cormat = np.zeros([3,3])
for i in range(3):
    for j in range(3):
        cormat[i,j] = stats.pearsonr(avg_speed[:,i], circ_conc[:,j])[0]
df = pd.DataFrame(cormat, index=['Trial 1 avg ind spd', 'Trial 2 avg ind spd', 'Trial 3 avg ind spd'],
                          columns=['Trial 1 circ conc',' Trial 2 circ conc','Trial 3 circ conc'])
df.to_csv('figures/avgspd_vs_circconc.csv')

# take a moving average with a 30-second window
time = np.arange(avg_speed.shape[0])/32/60
as_mov_avg = np.zeros(avg_speed.shape) + np.nan
cc_mov_avg = np.zeros(avg_speed.shape) + np.nan
radius = 15*32 # 15 seconds in either side
for i in range(avg_speed.shape[0]):
    if (i - radius >= 0) and (i + radius + 1 < avg_speed.shape[0]):
        as_mov_avg[i,:] = np.mean(avg_speed[i-radius:i+radius+1, :],axis=0)
        cc_mov_avg[i,:] = np.mean(circ_conc[i-radius:i+radius+1, :],axis=0)

# perform KPSS test with BOTH the raw data and the moving avg data
kpss_results = {'variable' : [], 'trial' : [], 'p<=' : [], '1% crit val' : [], 'statistic' : []}
kpss_settings = dict(regression='c', lags='auto')
for i in range(3):
    kpss_helper(
    timeseries = as_mov_avg[:,i],
    variable = 'avg spd mvg avg',
    trial = i + 1,
    kpss_results = kpss_results,
    nan_policy = 'exclude',
    **kpss_settings
    )

    kpss_helper(
    timeseries = cc_mov_avg[:,i],
    variable = 'circ conc mvg avg',
    trial = i + 1,
    kpss_results = kpss_results,
    nan_policy = 'exclude',
    **kpss_settings
    )

    kpss_helper(
    timeseries = avg_speed[:,i],
    variable = 'avg spd',
    trial = i + 1,
    kpss_results = kpss_results,
    nan_policy = 'strict',
    **kpss_settings
    )

    kpss_helper(
    timeseries = circ_conc[:,i],
    variable = 'circ conc',
    trial = i + 1,
    kpss_results = kpss_results,
    nan_policy = 'strict',
    **kpss_settings
    )
pd.DataFrame(kpss_results).to_csv('figures/kpss_results.csv', index=False)

# plot time series for circular concentration
for trial_num in range(3):
    plt.figure(figsize=[2.6,1.4])
    plt.title(f'trial {trial_num+1}')
    plt.plot(time, circ_conc[:,trial_num], 'k')
    plt.plot(time, cc_mov_avg[:,trial_num], 'r')
    plt.xlim([-0.5, 10.5])
    plt.ylim([-0.05,1.05])
    plt.xlabel('Time (min)', **font)
    plt.ylabel('Circular\nconcentration', **font)
    plt.savefig(f'figures/var_trial{trial_num+1}.pdf')
    plt.show()

# plot time series for average individual speed
for trial_num in range(3):
    plt.figure(figsize=[2.6,1.4])
    plt.title(f'trial {trial_num+1}')
    plt.plot(time, avg_speed[:,trial_num], 'k')
    plt.plot(time, as_mov_avg[:,trial_num], 'r')
    plt.xlim([-0.5, 10.5])
    plt.ylim([-0.5,7.5])
    plt.xlabel('Time (min)', **font)
    plt.ylabel('Avg ind spd\n(fish length / s)', **font)
    plt.savefig(f'figures/avg_speed_trial{trial_num+1}.pdf')
    plt.show()

plt.figure(figsize=[1,1.4])
plt.xlim([0.6, 0.65])
plt.ylim([2.5, 4.5])
plt.xlabel('Time-averaged\ncircular conc' , **font)
plt.ylabel('Time-averaged spd\n(fish length / s)', **font)
plt.plot(np.mean(circ_conc,axis=0), np.mean(avg_speed,axis=0), '.k')
rho, pval_2t = pearsonr(np.mean(circ_conc, axis=0), np.mean(avg_speed, axis=0))
assert rho > 0, "negative correlation, 1-tailed pval adjustment is invalid." # verify positive correlation
pval_1t = pval_2t/2
plt.text(0.625, 2.55, f'p={round(pval_1t,3)}', ha='center', va='bottom')
plt.savefig('figures/scatter.pdf', bbox_inches='tight')
plt.show()

# perform the test at different time series lengths using correlation coefficient
np.random.seed(1)
ts_len_grid = np.arange(20, 604, 20) # units of seconds; 600 seconds is 10 minutes
n_trials = 500
results = {'CC-perf' : np.zeros(ts_len_grid.size * n_trials),
           'spd-perf' : np.zeros(ts_len_grid.size * n_trials),
           'para' : np.zeros(ts_len_grid.size * n_trials),
           'pcorr' : np.zeros(ts_len_grid.size * n_trials),
           'ts_len' : np.zeros(ts_len_grid.size * n_trials)}
i = 0
for ts_len in ts_len_grid:
    for trial_idx in range(n_trials):
        max_start_idx = 600*32 - int(ts_len*32)
        if max_start_idx != 0:
            start_idx = np.random.choice(max_start_idx)
        else:
            start_idx = 0
        end_idx = start_idx + int(ts_len*32)
        results['ts_len'][i] = ts_len
        results['CC-perf'][i] = improved_permtest(avg_speed[start_idx:end_idx,:], circ_conc[start_idx:end_idx,:], corr_fxn=corrcoef)
        results['spd-perf'][i] = improved_permtest(circ_conc[start_idx:end_idx,:], avg_speed[start_idx:end_idx,:], corr_fxn=corrcoef)
        results['pcorr'][i] = pearsonr(np.mean(circ_conc[start_idx:end_idx,:],axis=0), np.mean(avg_speed[start_idx:end_idx,:],axis=0))[0] # divide pval by 2 to get 1-tailed pval
        results['para'][i] = pearsonr(np.mean(circ_conc[start_idx:end_idx,:],axis=0), np.mean(avg_speed[start_idx:end_idx,:],axis=0))[1]/2 # divide pval by 2 to get 1-tailed pval
        i += 1

df = pd.DataFrame(results)
df.loc[df['pcorr'] < 0, 'para'] = 1 # the parametric test was 1-tailed, looking for positive correlation

power = df.groupby('ts_len')[['CC-perf', 'spd-perf', 'para']].apply(lambda x: (x <= 0.05).mean())


plt.figure(figsize=[3,2])
power_colors = {
'para' : '#666666',
'CC-perf' : '#2bc4d6',
'spd-perf' : '#ffcc0e'
}
for condition in ['CC-perf', 'spd-perf', 'para']:
    plt.plot(power.index / 60, power[condition]*100, '.-', label=condition,
    color=power_colors[condition])
plt.xlabel('Segment length (min)', **font)
plt.ylabel('% segments detected', **font)
plt.ylim([-3, 103])
plt.legend()
plt.savefig('figures/detections.svg')
plt.show()

power.to_csv('figures/detections_data.csv')
```
