## Supplementary material for "Permute-match tests detect significant correlations between time series despite nonstationarity and limited replicates": Source Code 1: analyze_results_lite.py

```
import numpy as np
import pandas as pd
import matplotlib.pyplot as plt
from  matplotlib import rc, rcParams
from scipy.interpolate import interp1d
import os
import warnings

# matplotlib settings
plt.rcParams['svg.fonttype'] = 'none'
rcParams['axes.linewidth'] = 1.5
rcParams['xtick.major.width'] = 1.5
rcParams['xtick.major.size'] = 5
rcParams['ytick.major.width'] = 1.5
rcParams['ytick.major.size'] = 5

font = {'family' : 'Arial',
        'weight' : 'bold',
        'size'   : 12}

rc('font', **font)

line_specs = {}
line_specs['y_first'] = {'color':'#59a9ff', 'linestyle' : (0, (4, 6)), 'label' : 'sequential Y ▶ X'}
line_specs['x_first'] = {'color':'#ffcc00', 'linestyle' : (2, (4, 6)), 'label' : 'sequential X ▶ Y'}
line_specs['perm_only'] = {'color':'gray', 'linestyle' : '-', 'label' : 'permutation only'}
line_specs['simultaneous'] = {'color':'red', 'linestyle' : (4, (4, 6)), 'label' : 'simultaneous (X;Y)'}

curveplot_shape = [4.5, 1.12]
setups = ['perm_only', 'y_first', 'x_first', 'simultaneous']
# constant parameters
n_reps_grid = np.array([3,4,5])
significance_levels = np.array([0.05, 0.01, 0.001])
n_jobs = 200

# make outfolders
outfolder = ['figures', 'figures/05', 'figures/01', 'figures/001']
for folder in outfolder:
    if not os.path.exists(folder):
        os.makedirs(folder)

############################
##### Helper functions #####
############################
def get_power(data, significance_levels):
    """
    Args:
    * data (numpy array): data[i,j,k] is the p-value with the ith rep number,
      the jth parameter choice, and the kth simulation
    * significance_levels (numpy array): a list of significance levels

    Returns: numpy array `power` such that
    power[i,j,k] is the power with the ith rep number, the jth parameter choice,
    and the kth significance level
    """
    sig_size = significance_levels.size
    (rep_size, param_size, _) = data.shape
    power = np.zeros((rep_size, param_size, sig_size))
    for rep_idx in range(rep_size):
        for param_idx in range(param_size):
            for sig_idx in range(sig_size):
                alpha = significance_levels[sig_idx]
                power[rep_idx, param_idx, sig_idx] = np.mean(data[rep_idx, param_idx, :] <= alpha)
    return power

def check_for_file(fpath):
    if os.path.isfile(fpath):
        return True
    else:
        warnings.warn('No file found at: ' + fpath)
        return False

def make_figures(system_name, xlabel, xlim, ylim, xticks, yticks):
    # load data
    power = {}
    for setup in setups:
        data = [np.load(f'results/{setup}/{system_name}_{i+1}.npy') for i in range(n_jobs) if check_for_file(f'results/{setup}/{system_name}_{i+1}.npy')]
        data = np.concatenate(data, axis=2)
        param_grid = pd.read_csv(f'results/{setup}/{system_name}_legend.csv', index_col=0)
        # process data
        power[setup] = get_power(data, significance_levels)

    # plot results
    for sig_idx, sig_lvl in enumerate(significance_levels): # loop over significance levels
        plt.figure(figsize=curveplot_shape) # each significance level is a figure
        figure_data = {} # and each figure has an associated Excel file
        for repnum_idx, repnum in enumerate(n_reps_grid): # loop over replicate numbers
            plt.subplot(1, n_reps_grid.size, repnum_idx+1) # each replicate number is a panel
            subplot_data = {} # and each panel has an associated Excel tab
            plt.ylim(ylim)
            plt.xlim(xlim)
            plt.xticks(xticks)
            plt.xlabel(xlabel, **font)
            plt.title(f'{repnum} replicates', **font)
            if repnum_idx==0:
                plt.ylabel('Power', **font)
                plt.yticks(yticks)
            else:
                plt.yticks([])
            for setup in setups:
                plt.plot(param_grid.values.flatten(), power[setup][repnum_idx,:,sig_idx], **line_specs[setup])
                column_label = line_specs[setup]['label']
                subplot_data[column_label] = power[setup][repnum_idx,:,sig_idx]
            if sig_idx == 0 and repnum_idx == 0:
                plt.legend()
            figure_data[repnum] = pd.DataFrame(subplot_data, index=param_grid.values.flatten())

        plt.savefig(f'figures/{str(sig_lvl)[2:]}/{system_name}_pwr_curves.svg', bbox_inches='tight')
        with pd.ExcelWriter(f'figures/{str(sig_lvl)[2:]}/{system_name}_pwr_curves.xlsx') as writer:
            for repnum in n_reps_grid:
                figure_data[repnum].to_excel(writer, sheet_name=f'{repnum} replicates', index_label=xlabel)
        plt.close()

#######################################
##### asymmetric correlated noise #####
#######################################

system_name = 'asym_corr_noise'
xlabel = 'Cov($\epsilon_x$,$\epsilon_y$)'
xlim = [-0.05,0.55]
ylim = [-0.05,1.05]
xticks = [0, 0.25, 0.5]
yticks = [0, 0.5, 1]
make_figures(system_name, xlabel, xlim, ylim, xticks, yticks)

##########################
##### logistic (CCM) #####
##########################

system_name = 'logistic_ccm'
xlabel = '$r_{X,Y}$'
xlim = [-0.02,0.22]
ylim = [-0.05, 1.05]
xticks = [0, 0.1, 0.2]
yticks = [0, 0.5, 1]
make_figures(system_name, xlabel, xlim, ylim, xticks, yticks)

##########################
##### logistic (Pearson) #####
##########################

system_name = 'logistic_pcorr'
xlabel = '$r_{X,Y}$'
xlim = [-0.02,0.22]
ylim = [-0.05, 1.05]
xticks = [0, 0.1, 0.2]
yticks = [0, 0.5, 1]
make_figures(system_name, xlabel, xlim, ylim, xticks, yticks)
```
