## Supplementary material for "Permute-match tests detect significant correlations between time series despite nonstationarity and limited replicates": Source Code 1: benchmark_lite.py

```
from examples import *
from os.path import join
import pandas as pd
from copy import deepcopy

def param_scan(ground_truth_model, corr_fxn, varparam_grid, varparam_name,
               fixed_params, n_sims=1000, n_reps_grid=np.array([3,4,5]),
               data_length=100, correlation_params={},
               match_type='y-first'):
    """
    Args:
    * ground_truth_model (function): A function with the signature
        x, y = ground_truth_model(data_length=data_length, **params)
        where x and y are 1d numpy arrays.
    * corr_fxn (function): correlaiton statistic with the signature:
      K = corr_fxn(X,y)
      Here, K is a length m numpy array where K[i] is the correlaiton between y
      and the ith column of X.
    * varparam_grid (1d numpy array): values for the parameter that is to be varied
    * varparam_name (string): name of varied parameter; the name must be the same
      as the parameter name in the ground_truth_model.
    * fixed_params (dict): A dictionary of fixed parameters.
    * n_sims (int): number of simulations. Each simulation results in a single
      p-value.
    * n_reps (int): the number of replicates to use in testing
    * data_length (int): length of time series
    * correlation_params (dict): dictionary of params for corr_fxn
    * match_type (string): see improved_permtest for options.

    Returns:
    (pvals, legend)
    * results[i,j,k] is the pvalue on the kth simulation with
      n_reps_grid[i] replicates and cor_grid[j] correlation
    * legend is a Pandas data table indicating the scanned parameter values
    """
    results = np.zeros((n_reps_grid.size, varparam_grid.size, n_sims))
    for rep_idx in range(n_reps_grid.size):
        n_reps = n_reps_grid[rep_idx]
        for param_idx in range(varparam_grid.size):
            params = deepcopy(fixed_params)
            params[varparam_name] = varparam_grid[param_idx]
            results[rep_idx, param_idx, :] = benchmark(
                ground_truth_model=ground_truth_model, corr_fxn=corr_fxn,
                params=params, n_sims=n_sims, n_reps=n_reps,
                data_length=data_length, correlation_params=correlation_params,
                match_type=match_type)
    return results, pd.DataFrame({varparam_name : varparam_grid})

# Constants
seed = int(sys.argv[1])
n_sims = int(sys.argv[2])
"""
seed = 1
n_sims = 10
"""

n_reps_grid = np.array([3,4,5])
data_length = 100
setups = [
            {'match_type' : 'y-first', 'outpath' : 'results/y_first'},
            {'match_type' : 'x-first', 'outpath' : 'results/x_first'},
            {'match_type' : 'none', 'outpath' : 'results/perm_only'},
            {'match_type' : 'simultaneous', 'outpath' : 'results/simultaneous'}
          ]

#######################################
##### Correlated noise with trend #####
#######################################
# params
cor_grid = np.arange(0, 0.55, 0.01)
params = {'steepness' : 5}

# perform benchmark

for setup in setups:
    np.random.seed(seed)
    pvals, legend = param_scan(ground_truth_model=corrnoise_w_trend,
        corr_fxn=corrcoef, varparam_grid=cor_grid, varparam_name='cor',
        fixed_params=params, n_sims=n_sims, n_reps_grid=n_reps_grid,
        data_length=data_length, match_type=setup['match_type'])

    # save data & legend
    np.save(join(setup['outpath'],f'corr_noise_{seed}.npy'), pvals)
    if seed == 1:
        legend.to_csv(join(setup['outpath'],f'corr_noise_legend.csv'))

print('finished corr noise w trend!')
#############################################################
##### Correlated noise with trend, and different slopes #####
#############################################################
# params
cor_grid = np.arange(0, 0.55, 0.01)
params = {'x_steep_lim' : 2, 'y_steep_lim' : 5}

# perform benchmark

for setup in setups:
    np.random.seed(seed)
    pvals, legend = param_scan(ground_truth_model=corrnoise_w_trend,
        corr_fxn=corrcoef, varparam_grid=cor_grid, varparam_name='cor',
        fixed_params=params, n_sims=n_sims, n_reps_grid=n_reps_grid,
        data_length=data_length, match_type=setup['match_type'])

    # save data & legend
    np.save(join(setup['outpath'],f'asym_corr_noise_{seed}.npy'), pvals)
    if seed == 1:
        legend.to_csv(join(setup['outpath'],f'asym_corr_noise_legend.csv'))

print('finished corr noise w variable-slope trend!')
#######################################################
##### Logistic map with drifting parameters (CCM) #####
#######################################################
# params

def init_cond():
    return np.random.uniform(0.1, 0.9)
rxy_grid = np.arange(0,0.205,0.005)
params = {'x0' : init_cond,
          'y0' : init_cond,
          'rx' : np.linspace(3.72,3.82,data_length-1),
          'ry' : np.linspace(3.72,3.82,data_length-1)}

for setup in setups:
    np.random.seed(seed)
    # perform benchmark
    pvals, legend = param_scan(ground_truth_model=pairwise_logistic,
        corr_fxn=ccm, varparam_grid=rxy_grid, varparam_name='rxy',
        fixed_params=params, n_sims=n_sims, n_reps_grid=n_reps_grid,
        data_length=data_length, match_type=setup['match_type'])

    # save data & legend
    np.save(join(setup['outpath'],f'logistic_ccm_{seed}.npy'), pvals)
    if seed == 1:
        legend.to_csv(join(setup['outpath'],f'logistic_ccm_legend.csv'))

print('finished logistic w drifting parameters (ccm)!')


#######################################################################
##### Logistic map with drifting parameters (Pearson correlation) #####
#######################################################################
for setup in setups:
    np.random.seed(seed)
    # perform benchmark
    pvals, legend = param_scan(ground_truth_model=pairwise_logistic,
        corr_fxn=corrcoef, varparam_grid=rxy_grid, varparam_name='rxy',
        fixed_params=params, n_sims=n_sims, n_reps_grid=n_reps_grid,
        data_length=data_length, match_type=setup['match_type'])

    # save data & legend
    np.save(join(setup['outpath'],f'logistic_pcorr_{seed}.npy'), pvals)
    if seed == 1:
        legend.to_csv(join(setup['outpath'],f'logistic_pcorr_legend.csv'))

print('finished logistic w drifting parameters (pcorr)!')
```
