## Supplementary material for "Permute-match tests detect significant correlations between time series despite nonstationarity and limited replicates": Source Code 1: binomial_test.py

```
from scipy import stats
import pandas as pd
import numpy as np

n_sims = 5000
n_reps = 3
alpha = '05'
test_name = 'sequential Y ▶ X'
systems = ['asym_corr_noise', 'logistic_ccm']
coupling_label = {
    'asym_corr_noise' : 'Cov($\epsilon_x$,$\epsilon_y$)',
    'logistic_ccm' : '$r_{X,Y}$'
}

results = {
'system' : systems,
'# of replicates' : np.tile(n_reps, len(systems)),
'alpha' : np.tile(float(f'0.{alpha}'), len(systems)),
'test type' : np.tile(test_name, len(systems)),
'false positive rate' : [],
'guaranteed max FPR' : np.tile(1/n_reps**n_reps, len(systems)),
'binomial pvalue' : [],
'binomial pvalue (via cdf)' : [],
}

for system in systems:

    detection_rate = pd.read_excel(
        io=f'figures/{alpha}/{system}_pwr_curves.xlsx',
        sheet_name=f'{n_reps} replicates',
        header=0)
    cl = coupling_label[system]

    false_pos_rate = detection_rate.loc[detection_rate[cl]==0, test_name].values.item()

    assert (false_pos_rate * n_sims).is_integer(), "Expected the false positive rate times the number of simulations to be an integer!"

    pval_binomtest = stats.binomtest(
        k = int(false_pos_rate * n_sims),
        n = n_sims,
        p = 1 / (n_reps ** n_reps),
        alternative = 'less'
    ).pvalue

    pval_binom = stats.binom.cdf(
        k = int(false_pos_rate * n_sims),
        n = n_sims,
        p = 1 / (n_reps ** n_reps),
    )

    results['false positive rate'].append(false_pos_rate)
    results['binomial pvalue'].append(pval_binomtest)
    results['binomial pvalue (via cdf)'].append(pval_binom)

print(
    pd.DataFrame(results).set_index('system').T
)
```
