## Supplementary material for "Permute-match tests detect significant correlations between time series despite nonstationarity and limited replicates": Source Code 1: examples.py

```
import numpy as np
import sys
from mini_ccm import ccm_loocv, pcorr
from sklearn.linear_model import LinearRegression
from itertools import permutations
#### testing

def match_test(X, Y, corr_fxn, correlation_params={}, match_type='y-perfect'):
    """Check for an x-perfect or y-perfect match.

    Args:
    * X (2d numpy array): X is an nxm array, where n is the size of a sample and
      m is the number of samples
    * Y (2d numpy array): Y is an nxm array similar to X.
    * corr_fxn (function): correlaiton statistic with the signature:

      K = corr_fxn(X,y, **correlation_params)

      Here, K is a length m numpy array where K[i] is the correlation between y
      and the ith column of X.
    * correlation_params (dictionary): Parameters passed into `corr_fxn`
    * match_type (string): whether to check for a y-perfect match or an x-perfect
      match. A y-perfect match occurs if

      corr(X[:,i], Y[:,i]) > corr(X[:,j], Y[:,i]) for all i≠j.

      An x-perfect match occurs if

      corr(X[:,i], Y[:,i]) > corr(X[:,i], Y[:,j]) for all i≠j.

    Returns: True or False depending on whether the y-perfect or x-perfect match is found.

    Note: The asymmetric correlation signature enables certain optimizations in
    correlation functions, such as those that rely on KNN libraries where a single
    "y" library can be used repeatedly to compute correlations with many "x" partners
    """
    m = X.shape[1]
    M = np.zeros((m, m))
    for i in range(m):
        M[i,:] = corr_fxn(X, Y[:,i], **correlation_params) # M[i,j] is the corr between Y[i] and X[j], so each row is a y and each column is an x.

    if match_type == 'y-perfect':
        pass
    elif match_type == 'x-perfect':
        M = M.T
    else:
        raise ValueError('Options for match_type are `y-perfect` or `x-perfect`.')

    for i in range(m):
        row = M[i,:] # assuming y-perfect, extract ith row
        if not np.all(row[i] > row[np.arange(m)!=i]): # check whether the on-diag value is greater than off-diag values for this row
            return False
    return True

def permtest(X, Y, corr_fxn, correlation_params={}, n_perm='exact', verbose=False):
    """
    Args:
    * X (2d numpy array): X is an nxm array, where n is the size of a sample and
      m is the number of samples
    * Y (2d numpy array): Y is an nxm array similar to X.
    * corr_fxn (function): correlaiton statistic with the signature:

      K = corr_fxn(X,y)

      Here, K is a length m numpy array where K[i] is the correlaiton between y
      and the ith column of X.
    * n_perm (int): number of permutations to use in randomized permutation test
      Default is to perform an exact permutation test.
    * verbose (bool): If True, returns a tuple (pval, stat_rank), where pval is
      the p-value and stat_rank is the number of null correlations that are at
      or above the original correlations. Note that the original correlation is
      itself included in the set of null correlations. If False, returns
      stat_rank only.

    Returns stat rank (Verbose=False) or p-value and stat rank (verbose=True)
    """
    assert X.shape[1] == Y.shape[1]
    n = X.shape[1]
    M = np.zeros((n, n))
    for i in range(n):
        M[i,:] = corr_fxn(X, Y[:,i], **correlation_params)
        # M[i,j] is corr(Y[:,i], X[:,j])
    scores = np.zeros(np.math.factorial(n))
    stat = np.mean(M[np.arange(n),np.arange(n)])
    if n_perm == 'exact':
        perms = list(permutations(range(n)))
        nulldist = np.mean(M[perms,np.arange(n)],axis=1)
        stat_rank = np.sum(nulldist >= stat)
        pval = stat_rank / nulldist.size
    else:
        perms = [np.random.choice(n, size=n, replace=False) for _ in range(n_perm)]
        nulldist = np.mean(M[perms,np.arange(n)],axis=1)
        nulldist.append(stat)
        stat_rank = np.sum(nulldist >= stat)
        pval = stat_rank / nulldist.size
    if verbose:
        return pval, stat_rank
    else:
        return stat_rank

def improved_permtest(X, Y, corr_fxn, correlation_params={}, match_type='y-first'):
    """
    Args:
    * X (2d numpy array): X is an nxm array, where n is the size of a sample and
      m is the number of samples
    * Y (2d numpy array): Y is an nxm array similar to X.
    * corr_fxn (function): correlaiton statistic with the signature:

      K = corr_fxn(X,y)

      Here, K is a length m numpy array where K[i] is the correlaiton between y
      and the ith column of X.
    * correlation_params (dictionary): any parameters passed on to corr_fxn
    * match_type (string): type of match test. options are
        * 'y-first': listed as the sequential Y▶X test in the paper
        * 'x-first': listed as the sequential X▶Y test in the paper
        * 'simultaneous': listed as the simultaneous test in the paper
        * 'none': only a permutation test is performed

    Returns the p-value
    """
    p_perm, stat_rank = permtest(X, Y, corr_fxn, correlation_params=correlation_params, verbose=True)
    n_reps = Y.shape[1]
    if not (match_type in ['y-first', 'x-first', 'simultaneous', 'none']):
        raise ValueError("'match_type' must be one of the following: 'y-first', 'x-first', 'simultaneous', 'none'.")
    if stat_rank == 1:
        # y-first test
        if match_type == 'y-first':
            if match_test(X, Y, corr_fxn, correlation_params=correlation_params, match_type='y-perfect'):
                return 1 / (n_reps ** n_reps)
            elif match_test(X, Y, corr_fxn, correlation_params=correlation_params, match_type='x-perfect'):
                return 2 / (n_reps ** n_reps)

        # x-first test
        if match_type == 'x-first':
            if match_test(X, Y, corr_fxn, correlation_params=correlation_params, match_type='x-perfect'):
                return 1 / (n_reps ** n_reps)
            elif match_test(X, Y, corr_fxn, correlation_params=correlation_params, match_type='y-perfect'):
                return 2 / (n_reps ** n_reps)

        # simultaneous test
        if (match_type == 'simultaneous'):
            yperf = match_test(X, Y, corr_fxn, correlation_params=correlation_params, match_type='y-perfect')
            xperf = match_test(X, Y, corr_fxn, correlation_params=correlation_params, match_type='x-perfect')
            if yperf or xperf:
                return 2 / (n_reps ** n_reps)

        # permutation only
        if match_type == 'none':
            pass
    return p_perm

def corrcoef(X, y):
    data = np.zeros((X.shape[0], X.shape[1] + 1))
    data[:,0] = y
    data[:,1:] = X
    result = np.corrcoef(data, rowvar=False)
    return result[0,1:]

def ccm(X, y, embed_dim=2, score=pcorr):
    # X is putative causers; y is putative causee
    data = np.zeros([X.shape[0],X.shape[1]+1])
    data[:,0] = y
    data[:,1:] = X
    return ccm_loocv(data, embed_dim=embed_dim, score=score).score.values.astype(np.float)

#### ground truth models
# all ground truth models must have a data_length parameter
# ground truth models must return (x,y) where x and y are both 1d numpy arrays.

def noise_w_trend(eps, steep_lim, meas_noise_str, dist):
    data_length = eps.size
    if np.array(steep_lim).size == 1:
        steep = steep_lim
    else:
        lo, hi = steep_lim
        if dist == 'uniform':
            steep = np.random.uniform(lo, hi)
        elif dist == 'coinflip':
            steep = np.random.choice([lo, hi])
        else:
            raise ValueError('dist must be uniform or coinflip')
    trendline = np.arange(1, data_length+1) * steep / data_length
    meas_noise = np.random.normal(0, meas_noise_str, size=data_length)
    return eps + trendline + meas_noise

def corrnoise_w_trend(cor=0, data_length=100, x_steep_lim=(1,5), y_steep_lim=(5,5), meas_noise_x=0, meas_noise_y=0, dist='uniform', steepness=None):
    """
    Args:
      * cor (float): correlation, between 0 and 1
      * data length (int)
      * x_steep_lim (tuple or float): lower and upper limits of the steepness parameter for x. pass a float to set the steepness parameter directly
      * y_steep_lim (tuple or float): lower and upper limits of the steepness parameter for y. pass a float to set the steepness parameter directly
      * meas_noise_x (float): strength of measurement noise applied to x
      * meas_noise_y (float): strength of measurement noise applied to y
      * dist (string): distribution of steepness parameter. Options are 'uniform' or 'coinflip'.
      * steepness (None or float): if used, sets the steepness of both x and y, overriding x_steep_lim and y_steep_lim; optional parameter added for back-compatibility with earlier versions of the code

    Returns: a tuple (x,y) where x and y are correlated iid standard normal random variables, plus a linear time trend. For both x and y, the linear time trend is given by:

    t * steepness / data_length

    where t = 1,2, ...., data_length
    """
    C = np.array([[1, cor],
                  [cor, 1]])
    eps_x , eps_y = np.random.multivariate_normal([0,0], C, data_length).T
    if not (steepness is None):
        x_steep_lim = steepness
        y_steep_lim = steepness
    x = noise_w_trend(eps_x, x_steep_lim, meas_noise_x, dist)
    y = noise_w_trend(eps_y, y_steep_lim, meas_noise_y, dist)
    return x, y

def pairwise_logistic(x0, y0, rx, ry, rxy, data_length=100):
    """Generates data from the two-species logistic map with possibly variable r
        Args:
            x0 (float or function): initial condition for x
            y0 (float or function): initial condition for y
            data_length (int): length of time series
            rx (float or array): Length is data_length - 1.
            ry (float or array): Length is data_length - 1.
            rxy (float or array): effect of x on y. Length is data_length - 1.
        Returns:
            x and y from Logistic map process
    """
    try:
        rx[0]
    except:
        rx = np.zeros(data_length-1) + rx
    try:
        ry[0]
    except:
        ry = np.zeros(data_length-1) + ry
    try:
        rxy[0]
    except:
        rxy = np.zeros(data_length-1) + rxy
    result = np.zeros([data_length,2])
    if type(x0) is type(match_test):
        result[0,0] = x0()
    else:
        result[0,0] = x0
    if type(y0) is type(match_test):
        result[0,1] = y0()
    else:
        result[0,1] = y0
    for i in range(1, data_length):
        result[i,0] = result[i-1,0] * (rx[i-1] - rx[i-1] * result[i-1,0])
        result[i,1] = result[i-1,1] * (ry[i-1] - ry[i-1] * result[i-1,1] - rxy[i-1] * result[i-1,0])
    return result[:,0], result[:,1]

#### infrastructure
def benchmark(ground_truth_model, corr_fxn, params, n_sims=1000, n_reps=3,
              data_length=100, correlation_params={}, match_type='y-perfect'):
    """
    Args:
    * ground_truth_model (function): A function with the signature
        x, y = ground_truth_model(data_length=data_length, **params)
        where x and y are 1d numpy arrays.
    * corr_fxn (function): correlaiton statistic with the signature:
      K = corr_fxn(X,y)
      Here, K is a length m numpy array where K[i] is the correlaiton between y
      and the ith column of X.
    * params (dict): dictionary of params for ground_truth_model
    * n_sims (int): number of simulations. Each simulation results in a single
      p-value.
    * n_reps (int): the number of replicates to use in testing
    * data_length (int): length of time series
    * correlation_params (dict): dictionary of params for corr_fxn
    * match_type (string): type of match test. options are 'y-perfect', 'x-perfect', 'either', or 'none'. If 'either', then the perfect match test is Bonferroni-corrected. If 'none', then only a permutation test is performed.

    Returns an array of length n_sims whose ith entry is the p-value from the
    ith simulation
    """
    results = []
    for sim in range(n_sims):
        X = np.zeros([data_length, n_reps])
        Y = np.zeros([data_length, n_reps])
        for rep in range(n_reps):
            x,y = ground_truth_model(data_length=data_length, **params)
            X[:, rep], Y[:, rep] = x, y
        results.append(improved_permtest(X, Y, corr_fxn, correlation_params, match_type))
    return np.array(results)
```
