## Supplementary material for "Permute-match tests detect significant correlations between time series despite nonstationarity and limited replicates": Source Code 1: mini_ccm.py

```
import numpy as np
from scipy.stats import pearsonr
from scipy.spatial import distance_matrix
from pandas import DataFrame

def pcorr(x,y):
    return pearsonr(x,y)[0]

def ccm_loocv(data, embed_dim, tau=1, lib_sizes=[None], replace=False,
              n_replicates=1, weights='exp', pred_lag=0, score=pcorr,
              variable_libs=False):
    """
    Args:
        data (array): array with n rows and m+1 columns, where n is the number
            of time points and m is the number of putative causer variables. The
            first column is the putative causee
        embed_dim (int): embedding dimension
        tau (int): delay for delay embedding
        lib_sizes (iterable): list of library sizes. Default is maximum library size
        replace (False): replacement for libraries chosen from sizes below maximum
        n_replicates (int): number of replicate runs
        weights (string): weighting function. Choices are 'exp' or 'uniform'
        variable_libs (bool): If this parameter is True, and if lib_sizes is
            used, each delay vector uses a different random library.
    Returns:
        Pandas DataFrame. The column 'causer' denotes the index of the putative
            causer
    """
    X, _ = setup_problem(data, embed_dim=embed_dim, tau=tau, pred_lag=pred_lag)
    n_y = data.shape[-1] - 1
    Y = [setup_problem(data[:,[0,i+1]], embed_dim=embed_dim, tau=tau, pred_lag=pred_lag)[1] for i in range(n_y)]
    # STEP ONE: get true neigbor indices
    KX = distance_matrix(X,X)
    np.fill_diagonal(KX, np.inf) # not allowed to train on self
    (n, embed_dim) = X.shape
    RX = np.zeros_like(KX)
    for loc in range(n):
        RX[loc,:] = np.argsort(KX[loc])                # sort vec indices by dist to loc
    all_xnbrs = np.zeros([n, embed_dim+1])
    result = []
    for y_idx, y in enumerate(Y):
        for lib_size in lib_sizes:
            for rep in range(n_replicates):
                if lib_size is None:
                    all_xnbrs = RX[:, :embed_dim+1]
                else:
                    if variable_libs:
                        rnd_lib = my_random_choice(a=n, size=lib_size, n=n, replace=replace).T
                        for loc in range(n):
                            mask = np.in1d(RX[loc,:], rnd_lib[loc,:])
                            all_xnbrs[loc,:] = RX[loc,:][mask][:embed_dim+1]
                    else:
                        rnd_lib = np.random.choice(n, size=lib_size, replace=replace)
                        for loc in range(n):
                            mask = np.in1d(RX[loc,:], rnd_lib)
                            all_xnbrs[loc,:] = RX[loc,:][mask][:embed_dim+1]
                # STEP TWO: compute observed cross-map skill
                if weights == 'uniform':
                    yhat = np.mean(y[all_xnbrs.astype(np.int)], axis=1) # compute observed xmap skill
                elif weights == 'exp':
                    yhat = [np.sum( y[all_xnbrs[loc,:].astype(np.int)] *
                                    get_weights(KX[loc,all_xnbrs[loc,:].astype(np.int)])
                                  )
                            for loc in range(n)]
                    yhat = np.array(yhat)
                else:
                    raise ValueError('weights must be `exp` or `uniform`.')
                result.append([lib_size, y_idx, score(y,yhat)])
    result = np.array(result)
    return DataFrame(result, columns=['lib_size', 'causer', 'score'])

def get_weights(distances):
    """Performs weighting in KNN for Simplex projection

    Args:
      * distances_ (array) Array of distances

    Returns:
      * weights
    """
    distances = np.array(distances, copy=True).astype(np.float)
    if len(distances.shape) == 1:
        distances = distances.reshape(1,-1)
    if np.any(distances == 0): # deal with zeros
        distances[distances != 0] = np.inf
        distances[distances == 0] = 1
    min_dist = np.min(distances, axis=1).reshape(-1,1)
    u = np.exp(-distances / min_dist)
    weights = u / np.sum(u, axis=1).reshape(-1,1)
    return weights

def my_random_choice(a, size, n, replace=False):
    """
    Args:
        a (int): number of items to choose from
        size (int): number of items to choose on each replicate trial
        n (int): number of trials
        replace (bool): choose with or without replacement

    Returns:
        an (size x n) matrix where each column is a trial
    """
    if replace:
        return np.random.choice(a, size=[size, n], replace=True)
    else:
        return np.argsort(np.random.random([a, n]), axis=0)[:size, :]

def setup_problem(data, embed_dim, tau, pred_lag=0):
    """Prepares a "standard matrix problem" from the time series data

    Args:
        data (array): A 2d array with two columns where the first column
            will be used to generate features and the second column will
            be used to generate the response variable
        embed_dim (int): embedding dimension (delay vector length)
        tau (int): delay between values
        pred_lag (int): prediction lag
    """
    x = data[:,0]
    y = data[:,1]
    feat = []
    resp = []
    idx_template = np.array([pred_lag] + [-i*tau for i in range(embed_dim)])
    for i in range(x.size):
        if np.min(idx_template + i) >= 0 and np.max(idx_template + i) < x.size:
            feat.append(x[idx_template[1:] + i])
            resp.append(y[idx_template[0] + i])
    return np.array(feat), np.array(resp)

def choose_embed_params(ts, ed_grid=np.arange(1,9), tau_grid=np.arange(1,9), weights='exp', score=pcorr):
    data = np.zeros([len(ts), 2])
    data[:,0] = ts
    data[:,1] = ts
    result = []
    for embed_dim in ed_grid:
        for tau in tau_grid:
            rho = ccm_loocv(data, embed_dim=embed_dim, tau=tau, pred_lag=1,
                                 weights=weights, score=score).score.values.item()
            result.append([embed_dim, tau, rho])
    result = np.array(result)
    winner_idx = np.argmax(result[:,2])
    return result[winner_idx, 0].astype(np.int), result[winner_idx, 1].astype(np.int)
```
