## Supplementary material for "Permute-match tests detect significant correlations between time series despite nonstationarity and limited replicates": Source Code 1: plot_example_timeseries.py

```
import matplotlib.pyplot as plt
from  matplotlib import rc, rcParams
from copy import deepcopy
from examples import *
import os

# matplotlib settings
plt.rcParams['svg.fonttype'] = 'none'
rcParams['axes.linewidth'] = 1.5
rcParams['xtick.major.width'] = 1.5
rcParams['xtick.major.size'] = 5
rcParams['ytick.major.width'] = 1.5
rcParams['ytick.major.size'] = 5

font = {'family' : 'Arial',
        'weight' : 'bold',
        'size'   : 12}

rc('font', **font)

# make outfolders
outfolder = ['figures', 'figures/example_time_series']
for folder in outfolder:
    if not os.path.exists(folder):
        os.makedirs(folder)

# constants
data_length = 100
np.random.seed(1)

############################
##### correlated noise #####
############################

params = {'steepness' : 5}
x,y = corrnoise_w_trend(cor=0, **params)

"""
plt.figure(figsize=[1.8,1.2])
plt.plot(np.arange(1,101), x, 'k', linewidth=1)
plt.plot(np.arange(1,101), y, 'r', linewidth=1)
plt.xlabel('time', **font)
plt.ylim([-2.5,7.5])
plt.xlim([0,100])
plt.title('Cov($\epsilon_x$,$\epsilon_y$) = 0', **font)
plt.savefig('figures/example_time_series/corr_noise_indep.svg')
plt.show()

x,y = corrnoise_w_trend(cor=0.3, **params)
plt.figure(figsize=[1.8,1.2])
plt.plot(np.arange(1,101), x, 'k', linewidth=1)
plt.plot(np.arange(1,101), y, 'r', linewidth=1)
plt.xlabel('time', **font)
plt.ylim([-2.5,7.5])
plt.xlim([0,100])
plt.title('Cov($\epsilon_x$,$\epsilon_y$) = 0.3', **font)
plt.savefig('figures/example_time_series/corr_noise_dep.svg')
plt.show()
"""

#######################################
##### asymmetric correlated noise #####
#######################################

params = {'x_steep_lim' : 2, 'y_steep_lim' : 5}
x,y = corrnoise_w_trend(cor=0, **params)

plt.figure(figsize=[1.8,1.2])
plt.plot(np.arange(1,101), x, 'k', linewidth=1)
plt.plot(np.arange(1,101), y, 'r', linewidth=1)
plt.xlabel('time', **font)
plt.ylim([-2.5,7.5])
plt.xlim([0,100])
plt.title('Cov($\epsilon_x$,$\epsilon_y$) = 0', **font)
plt.savefig('figures/example_time_series/asym_corr_noise_indep.svg')
plt.show()

x,y = corrnoise_w_trend(cor=0.3, **params)
plt.figure(figsize=[1.8,1.2])
plt.plot(np.arange(1,101), x, 'k', linewidth=1)
plt.plot(np.arange(1,101), y, 'r', linewidth=1)
plt.xlabel('time', **font)
plt.ylim([-2.5,7.5])
plt.xlim([0,100])
plt.title('Cov($\epsilon_x$,$\epsilon_y$) = 0.3', **font)
plt.savefig('figures/example_time_series/asym_corr_noise_dep.svg')
plt.show()


########################
##### Logistic map #####
########################

def init_cond():
    return np.random.uniform(0.1, 0.9)
rxy_grid = np.arange(0,0.205,0.005)
params = {'x0' : init_cond,
          'y0' : init_cond,
          'rx' : np.linspace(3.72,3.82,data_length-1),
          'ry' : np.linspace(3.72,3.82,data_length-1)}

x,y = pairwise_logistic(rxy=0, **params)
plt.figure(figsize=[1.8,1.2])
plt.subplot(2,1,1)
plt.plot(np.arange(1,101), x, 'k', linewidth=1)
plt.ylabel('$x$', rotation=0)
plt.ylim([-0.1,1.1])
plt.xticks([])
plt.xlim([0,100])
plt.title('$r_{xy}$ = 0', **font)
plt.subplot(2,1,2)
plt.plot(np.arange(1,101), y, 'r', linewidth=1)
plt.ylabel('$y$', rotation=0)
plt.ylim([-0.1,1.1])
plt.xlabel('time', **font)
plt.xlim([0,100])
plt.savefig('figures/example_time_series/logistic_indep.svg')
plt.show()

# poincare plot
np.random.seed(999)
plt.figure(figsize=[2, 1.8])
for i in range(10):
    x,y = pairwise_logistic(rxy=0, **params)
    sc = plt.scatter(x[:-1], x[1:], 5, c=np.arange(1,100), cmap='cool')
plt.plot([0.4, 0.4 , 0.6,  0.6, 0.4 ],
         [0.9, 0.96, 0.96, 0.9, 0.9], 'k')
plt.colorbar(sc, ticks=[1,99])
plt.xticks([0.2, 0.6, 1], **font)
plt.yticks([0.2, 0.6, 1], **font)
plt.xlabel('$x_{t}$', **font)
plt.ylabel('$x_{t+1}$', **font, rotation=0)
plt.savefig('figures/example_time_series/logistic_poincare.svg')
plt.show()

np.random.seed(999)
plt.figure(figsize=[0.8, 0.8])
for i in range(10):
    x,y = pairwise_logistic(rxy=0, **params)
    plt.scatter(x[:-1], x[1:], 5, c=np.arange(1,100), cmap='cool')
plt.xlim([0.4, 0.6])
plt.ylim([0.9, 0.96])
plt.xticks([])
plt.yticks([])
plt.savefig('figures/example_time_series/logistic_poincare_insert.svg')
plt.show()

x,y = pairwise_logistic(rxy=0.2, **params)
plt.figure(figsize=[1.8,1.2])
plt.subplot(2,1,1)
plt.plot(np.arange(1,101), x, 'k', linewidth=1)
plt.ylabel('$x$', rotation=0)
plt.ylim([-0.1,1.1])
plt.xticks([])
plt.xlim([0,100])
plt.title('$r_{xy}$ = 0.2', **font)
plt.subplot(2,1,2)
plt.plot(np.arange(1,101), y, 'r', linewidth=1)
plt.ylabel('$y$', rotation=0)
plt.ylim([-0.1,1.1])
plt.xlabel('time', **font)
plt.xlim([0,100])
plt.savefig('figures/example_time_series/logistic_dep.svg')
plt.show()
```
