## Supplementary material for "Permute-match tests detect significant correlations between time series despite nonstationarity and limited replicates": Source Code 1: run_lite.sh

```
#$ -l tmem=6G
#$ -l h_vmem=6G
#$ -l h_rt=24:0:0 # at most 25 hours runtime
#$ -S /bin/bash
#$ -j y
#$ -N bench-lite
#$ -t 1-200
#$ -o /PATH/TO/CODE/permute_match_code/main_benchmark/logs # REPLACE WITH AN ACTUAL LOG PATH
cd /PATH/TO/CODE/permute_match_code/main_benchmark # REPLACE WITH ACTUAL PATH TO THE FOLDER
hostname
date
python3 benchmark_lite.py $SGE_TASK_ID 25
date
```
