## Supplementary material for "Permute-match tests detect significant correlations between time series despite nonstationarity and limited replicates": Source Code 1: readme.md

### *Permute-match tests detect significant correlations between time series despite nonstationarity and limited replicates*

This readme file describes the libraries used for the code and instructions to reproduce the figures.

### Dependencies
The main libraries used in this code are numpy, pandas, seaborn, matplotlib, scipy, statsmodels, and scikit-learn. For completeness, a full list of packages and versions is provided for the two machine setups where code was run. The file `benchmark_lite.py` was run on a computing cluster with the environment specified at the file `environments/server.txt` and all other scripts were run in the environment specified at the file `environments/local.txt` and `environments/local.yml`.

### Example Time Series: Fig 2C, S2B, and S2C
To reproduce these figures, navigate to `main_benchmark/` and run `python plot_example_timeseries.py`. This will generate several files in `main_benchmark/figures/example_time_series/`. In particular, Fig 2C is `asym_corr_noise_dep.svg`, Fig S2B is  `logistic_dep.svg` and Fig S2C is made from `logistic_poincare.svg` and `logistic_poincare_insert.svg`.

### Benchmark Results: Fig 2D, S2D, and S3
Navigate to `main_benchmark/` and run the command

`python benchmark_lite.py $job_id 25`

in a loop where `$job_id` runs from 1 to 200. The file `run_lite.sh` was what I used with an SGE job scheduler to do this in parallel, but in principle you could do it in serial on a single computer. Then, run `analyze_results_lite.py` to generate the results. They will appear in the folders `figures/$alpha` where `$alpha` is 05, 01, or 001, corresponding to the figures with the significance levels of 0.05, 0.01, and 0.001.

Lastly, you may run `binomial_test.py` to print out the results of two exact binomial tests that have in-text citations in the section "The permute-match test’s false positive rate depends on the process tested".

### Real Data Example: Fig 3
For this you will need to download the data from https://drive.google.com/drive/folders/1UmzlX-yJhzQ5KX5rGry8wZgXvcz6HefD and save the folder on your computer. Then go into the script `fish_example/fish_example.py` and change the file path at line 54 from the file path on my computer to the file path where the downloaded folder is stored on your computer. Then run the program `fish_example/fish_example.py` and this will generate the figures and results in the folder `fish_example/figures`. Note: The alignment variable, denoted "mean resultant length" in the mauscript, is denoted "circ conc" or CC in the output files.

### Runtime benchmark: Source data 1
Navigate to `runtime/` and run the command `python runtime.py`. Results will appear in the `runtime/` folder.
