## Supplementary material for "Permute-match tests detect significant correlations between time series despite nonstationarity and limited replicates": Source Code 1: runtime.py

```
import sys
sys.path.append('../main_benchmark')
from examples import *
import timeit
import pandas as pd
import matplotlib.pyplot as plt
from matplotlib.pyplot import rcParams, rc
import time

plt.rcParams['svg.fonttype'] = 'none'
rcParams['axes.linewidth'] = 1.5
rcParams['xtick.major.width'] = 1.5
rcParams['xtick.major.size'] = 5
rcParams['ytick.major.width'] = 1.5
rcParams['ytick.major.size'] = 5

font = {'family' : 'Arial',
        'weight' : 'bold',
        'size'   : 12}

rc('font', **font)

ground_truth_model = corrnoise_w_trend
corr_fxn = corrcoef
params = {'x_steep_lim' : 2, 'y_steep_lim' : 5, 'cor' : 0.2}
data_length = 100
match_type = 'x-first'
correlation_params = {}

##### Get runtime estimates via timeit
n_reps_grid = [3, 4, 5, 6, 7, 8, 9, 10]
n_sims_grid = [50000, 20000, 10000, 10000, 2000, 500, 100, 10]
runtimes = []
for i in range(len(n_reps_grid)):
    n_reps = n_reps_grid[i]
    n_sims = n_sims_grid[i]
    X = np.zeros([data_length, n_reps])
    Y = np.zeros([data_length, n_reps])
    np.random.seed(1)
    for rep in range(n_reps):
        x,y = ground_truth_model(data_length=data_length, **params)
        X[:, rep], Y[:, rep] = x, y
    rt = timeit.timeit(stmt='improved_permtest(X, Y, corr_fxn, correlation_params, match_type)', globals=globals(), number=n_sims)
    runtimes.append(rt)
df = pd.DataFrame({'reps' : n_reps_grid,
                   'sims' : n_sims_grid,
                   'total_time' : runtimes})
df['time_per_sim'] = df['total_time'] / df['sims']
df.to_csv('runtime.csv')
plt.figure(figsize=[4, 3])
plt.semilogy(df.reps, df.time_per_sim, 'o-k')
plt.yticks([10**-3, 10**-2, 10**-1, 1, 10], **font)
plt.xticks(np.arange(3, 11), **font)
plt.xlabel('Number of replicates', **font)
plt.ylabel('Time (seconds)', **font)
plt.savefig('runtime.svg')
plt.close()

# do it again with naive approach: uses a simpler timer but
# benefits from more random systems

runtimes = []
for i in range(len(n_reps_grid)):
    np.random.seed(1)
    n_reps = n_reps_grid[i]
    n_sims = n_sims_grid[i]
    X = np.zeros([data_length, n_reps, n_sims])
    Y = np.zeros([data_length, n_reps, n_sims])
    for sim in range(n_sims):
        for rep in range(n_reps):
            x,y = ground_truth_model(data_length=data_length, **params)
            X[:, rep, sim], Y[:, rep, sim] = x, y
    results = []
    start_time = time.time()
    for sim in range(n_sims):
        results.append(improved_permtest(X[:,:,sim], Y[:,:,sim], corr_fxn, correlation_params, match_type))
    end_time = time.time()
    runtimes.append(end_time - start_time)

df2 = pd.DataFrame({'reps' : n_reps_grid,
                   'sims' : n_sims_grid,
                   'total_time' : runtimes})
df2['time_per_sim'] = df2['total_time'] / df2['sims']

plt.semilogy(df.reps, df.time_per_sim, '-')
plt.semilogy(df2.reps, df2.time_per_sim, ':')
plt.savefig('runtime_compare.svg')
```
